# *Salmonella* subverts autophagy balancing bacterial fate and cellular inflammation

**DOI:** 10.1101/2020.11.15.383372

**Authors:** Chak Hon Luk, Wei Yu, Ludovic Deriano, Jost Enninga

**Affiliations:** Dynamics of Host-Pathogen Interactions Unit, Institut Pasteur, UMR3691 CNRS, Paris 75015, France; Université de Paris, Sorbonne Paris Cité, Paris, France; Genome Integrity, Immunity and Cancer Unit, Equipe Labellisée Ligue Contre le Cancer, Department of Immunology, Department of Genomes and Genetics, Institut Pasteur, 75015 Paris, France

**Keywords:** *Salmonella* Typhimurium, *Salmonella* lifestyle, vacuolar rupture, autophagy, host-pathogen interaction

## Abstract

*Salmonella* Typhimurium (*S.* Typhimurium) is an enteric bacterium capable of invading a wide range of host cell types and adopting different intracellular lifestyles for survival. Host endocytic trafficking and autophagy have been implied to regulate the *S.* Typhimurium subcellular localization and survival. To reveal alternative host regulators on *S.* Typhimurium lifestyle, we combined a novel fluorescent reporter, *Salmonella* Intracellular Analyzer (SINA) with haploid forward genetic screening. This identified transcription factor c-MYC as a negative regulator of *S.* Typhimurium cytosolic lifestyle via stabilizing the *Salmonella*-containing vacuole (SCV). We further confirmed that c-MYC downstream regulated LC3 acts to maintain SCV stability and limits *S.* Typhimurium cytosolic lifestyle. We demonstrated that LC3 is recruited to the SCV prior to the endomembrane damage marker Galectin 3, and it regulates SCV stability independent of the autophagosome adaptor NDP52. The LC3 processing enzymes ATG3 and ATG4 reciprocally act on SCV stability, where the loss of LC3-PE conjugation in the absence of ATG3 limits SCV damages. We further identified the dosage-dependent function of the *S.* Typhimurium effector SopF in mediating SCV stability by actively avoiding LC3 recruitment to the proximity of the SCV to reduce its catastrophic rupture and host cell death. Altogether, we offer insights on the significance of cellular transcription profile in the determination of *S.* Typhimurium pathophysiology as well as the underlying host-evasion strategy of *S.* Typhimurium.

## Introduction

*Salmonella enterica* serovar Typhimurium (*S.* Typhimurium) is an enteric bacterial pathogen, being one of the major causative agents for global food-borne illnesses. *S.* Typhimurium could be detected in natural reservoirs from the environment as well as animal farming, food manufacturing production lines, and be prevalent among patients in clinical settings (Stanaway et al., 2019). In the majority of the *S.* Typhimurium-caused illnesses, the pathogens were transmitted to humans via ingestion of contaminated food or water sources, where host invasion commences at the human gastrointestinal tract.

Surviving the acidic pH of the gastric acid, *S.* Typhimurium reach the host intestine, the first susceptible host tissue against *S.* Typhimurium invasion. Among the luminal *S.* Typhimurium, a fraction of *S.* Typhimurium expresses the *Salmonella* Pathogenicity Island 1 (SPI-1)-encoded Type III Secretion System 1 (T3SS1) and its cognate effectors to actively induce bacterial uptake by the non-phagocytic epithelial cells (Hume et al., 2017). Upon entry into enterocytes, *S.* Typhimurium are encapsulated in an endocytic compartment coined *Salmonella*-containing vacuole (SCV) that stands at the crossroad of maturation by remodeling or rupture. In 80-90% of the SCV formed, *S.* Typhimurium express the *Salmonella* Pathogenicity Island 2 (SPI-2)-encoded Type III Secretion System 2 (T3SS2) and effectors to halt the lysosomal fusion and remodel the SCV into a viable niche (Jennings et al., 2017; Lou et al., 2019). Shortly upon SCV formation following *S.* Typhimurium uptake, the SCV dynamically interacts with surrounding macropinosomes mediated by host SNAP receptor (SNARE) proteins and *S.* Typhimurium effector SopB, which control the SCV stability (Fredlund et al., 2018; Stévenin et al., 2019). In parallel, the host autophagy pathway has also been implied in the SCV stability maintenance, where the disruption of autophagy leads to elevated SCV rupture and reduced *S*. Typhimurium intracellular survival (Birmingham et al., 2006; Kreibich et al., 2015; Yu et al., 2014). Recently, the *S.* Typhimurium effector SopF was reported to maintain SCV integrity and acts on the autophagy system in the context of vacuolar rupture (Lau et al., 2019; Xu et al., 2019). This is relevant as *S.* Typhimurium released into the host cytosol are first challenged by autophagy degradation, where the escaped *S*. Typhimurium undertake a fast-growing phenotype, coined hyper replication (Knodler et al., 2010; Wu et al., 2020). The hyper replicating *S.* Typhimurium in the host cytosol activates the host non-canonical inflammosome, leading to host cell extrusion and pyroptotic death, which releases the invasion-primed hyper replicating *S.* Typhimurium into the gut lumen. This phenomenon has been proposed to support rapid *S.* Typhimurium spreading and tissue colonization in the host gut (Knodler et al., 2014; Sellin et al., 2014). Despite the extensive characterization of the different *S.* Typhimurium intracellular phenotypes and the recognition of the involvement of intracellular trafficking pathways in controlling these phenotypes, alternative host regulators remain largely unexplored.

In this work, we combined our recently developed *<u>S</u>almonella* <u>IN</u>tracellular <u>A</u>nalyzer (SINA) system with a haploid cell screening platform, we determined the roles of c-MYC and LC3 in restricting *S.* Typhimurium cytosolic via SCV stabilization. With fluorescent microscopy and flow cytometry, we determined the significance of LC3-PE conjugation on SCV catastrophic damage. Ultimately, we identified that the role of *S.* Typhimurium effector SopF in SCV maintenance is mediated by the avoidance of LC3 conjugation on SCV, serving a function to control the fate of infected host cells.

## Results

### GeneTrap Screening for host regulators of *S.* Typhimurium lifestyles

The distinct intracellular lifestyles of *S.* Typhimurium in epithelial cells have been reported, which exhibit unique subcellular localization and replication rate. With the aid of our recently developed multiplex fluorescent reporter, *Salmonella* Intracellular Analyzer (SINA), the intracellular lifestyle of *S.* Typhimurium can be precisely depicted at single-cell and single-bacterium levels (Figure S1) (unpublished). To date, a number of host endocytic trafficking pathways as well as bacterial effectors have been proposed to alter the SCV stability and subsequent *S.* Typhimurium hyper replication (Fredlund et al., 2018; Kreibich et al., 2015; Lau et al., 2019; Santos et al., 2015; Stévenin et al., 2019; Wrande et al., 2016; Xu et al., 2019; Yu et al., 2014). Despite extensive efforts in the recent years, a functional screen on host factors for *S.* Typhimurium lifestyle regulator remains lacking. To identify host factors that regulate *S.* Typhimurium hyper replication, we combined our SINA reporter system with a retrovirus-mediated monoallelic cell screening approach. The screen approach employed a haploid cell line, eHAP, which is randomly mutagenized by the transduction of retrovirus harboring a Gene Trap cassette (Carette et al., 2009). The pool of mutagenized cells was then expanded and infected with *S.* Typhimurium harboring SINA1.2. Individual pools of infected cells harboring ≥ 2-fold increase of cytosolic *S.* Typhimurium as compared to the non-mutagenized control were selected, cells harboring cytosolic *S.* Typhimurium were enriched by cell sorting. The insertion sites were mapped using non-restriction linear amplification PCR (nrLAM-PCR) and next-generation sequencing, the obtained reads were aligned to the human genome to identify the screen candidates (Figure 1A) (Paruzynski et al., 2010). For the screen, *S.* Typhimurium harbored SINA1.2, which constitutes of a constitutively expressed DsRed and cytosolic-responsive smURFP (Figure 1B). This variant of SINA enabled the selection of cells that are infected (DsRed^+^) and harboring cytosolic *S.* Typhimurium (smURFP^+^) (Figure S2). As eHAP is an engineered cell line derived from chronic leukemia cell KBM7, we first verified the invasiveness of *S.* Typhimurium in eHAP and the adoption of cytosolic lifestyle within (Figure 1C) (Essletzbichler et al., 2014). In non-mutagenized eHAP, cytosolic *S.* Typhimurium is present in 0.06% of infected cells, we confirmed the incidence of cytosolic *S.* Typhimurium could be boosted with the amiloride treatment (Stévenin et al., 2019). In mutagenized eHAP, most individual pools exhibited a comparable incidence of cytosolic *S.* Typhimurium (~0.05%), while pools selected for cell sorting harbors ≥ 2-fold increase of cytosolic *S.* Typhimurium (Figure 1D). Sequencing libraries were prepared to capture the insertion sites of the Gene Trap cassette, and subsequently mapped on the human genome (Tables S1 and S2, Figure S3).

**Figure 1.**
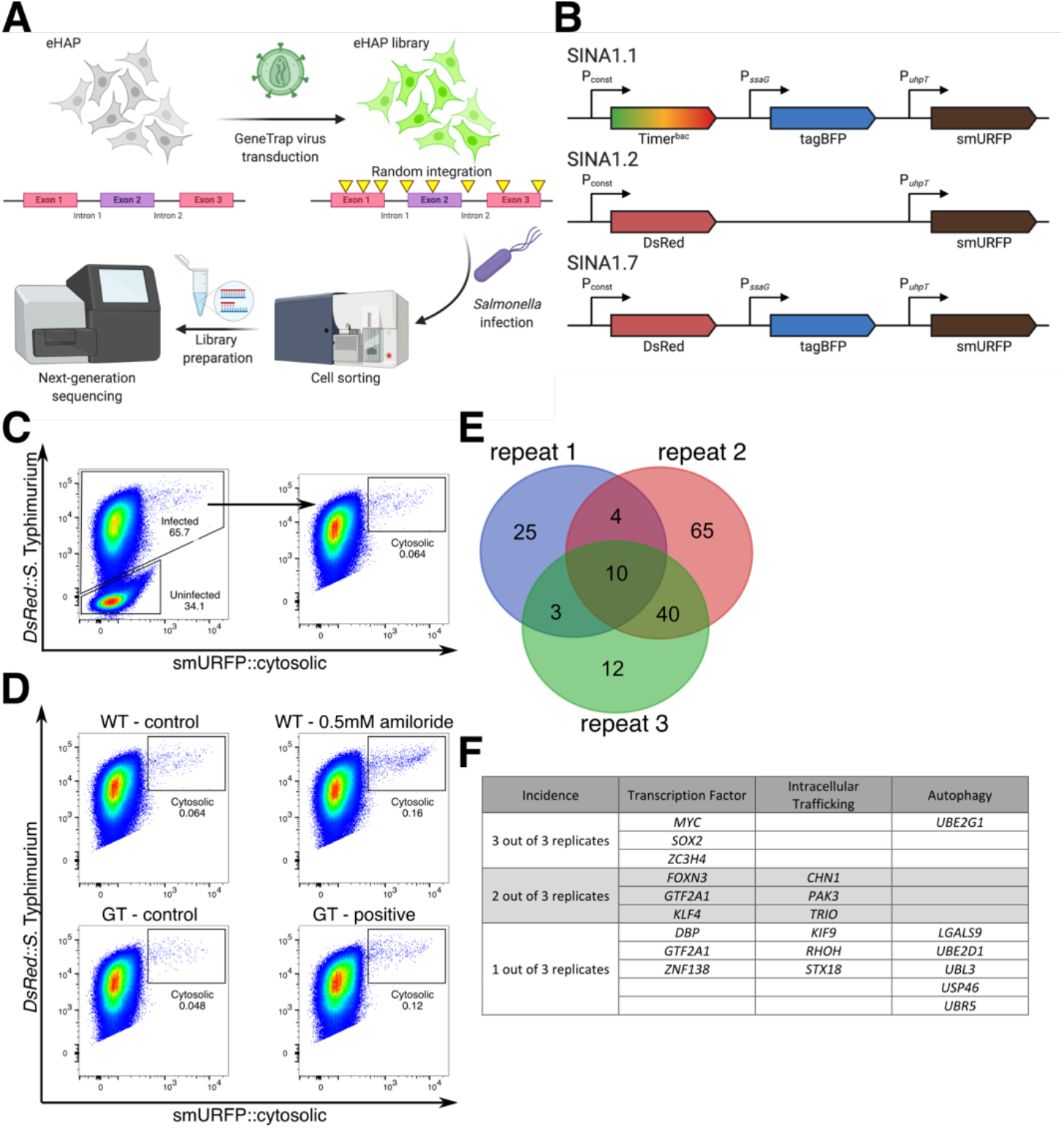
Genetic screen on host regulators on *S.* Typhimurium cytosolic lifestyle by combining SINA system and monoallelic cell screening approach. (A) Schematic illustration of the workflow of monoallelic cell screening approach, outlining the mutagenesis of eHAP cells, *S.* Typhimurium infection, cell enrichment, library preparation and sequencing. (B) Schematic diagrams of variants of SINA system. SINA1.1 was used for simultaneous profiling of *S.* Typhimurium localization and replication rate; SINA1.2 was used for eHAP infection; SINA1.7 was used for profiling of *S.* Typhimurium localization in conjunction with transgene ectopic expression in host cells. (C) Gating strategy of eHAP cells infected by SINA1.2-harboring *S.* Typhimurium, where infected cells, and then infected cells with cytosolic *S.* Typhimurium were gated. (D) Representative flow cytometry plots of SINA1.2-harboring *S.* Typhimurium infected wildtype eHAP (Top Left), wildtype eHAP treated with 0.5 mM amiloride (Top Right), mutagenized eHAP not subjected to enrichment (Bottom Left) and mutagenized eHAP subjected to enrichment (Bottom Right). (E) Venn diagram depicting the filtered hits obtained in triplicated experiments. (F) Partial list of the enriched host genes identified in the mutagenized eHAP cells enriched by cell sorting.

From the identified candidates, we first selected candidates with > 50 reads per library to obtain 47 candidates with incidence in 2 out of 3 repeats and 10 candidates with incidence in all 3 repeats (Figure 1E). We considered the 57 hits and removed the candidates that did not show a ≥ 2-fold increase as compared to the unenriched pool, ultimately yielding a total of 45 candidates (Table S1). Among the candidates, we have identified a series of host factors involved in transcription regulation, intracellular trafficking and autophagy (Figure 1F). Among the transcription factors, we observed three of the four Yamanaka factors, which hold essential functions in inducing and regulating cellular stemness (Takahashi and Yamanaka, 2006).

### c-MYC restricts *S.* Typhimurium cytosolic lifestyle independent of cell cycle

With the three identified Yamanaka factors, c-MYC, KLF4 and SOX2, c-MYC has been implied in regulating anti-mycobacterial responses mediated by the mitogen-activated protein kinases (MAPK)/ extracellular signal-regulated kinases (ERK) pathway and regulate autophagosome and lysosome formation (Annunziata et al., 2019; Toh et al., 2013; Yim et al., 2011). To verify the functional role of the identified candidates, we individually knocked down the candidates and determine the level of *S.* Typhimurium hyper replication. At 6 h pi, we observed a significant increase of hyper replication in c-MYC knockdown but not in KLF4 and SOX2 knockdown (Figures 2A and S4) To further understand whether c-MYC regulates cytosolic lifestyle via SCV integrity maintenance or autophagy, we determined the incidence of SCV rupture by quantifying the recruitment of Galectin 3 towards *S.* Typhimurium proximity using fluorescent microscopy. Galectin 3 is a member of the beta-galactoside-binding proteins family that has been established as a marker for vacuole lysis (Paz et al., 2010). We observed an increase SCV rupture in c-MYC knockdown, suggesting c-MYC limits cytosolic lifestyle via SCV integrity maintenance (Figure 2B). To further confirm the role of c-MYC in limiting *S.* Typhimurium cytosolic lifestyle, we used c-MYC inhibitor 10074-G5 that inhibits DNA binding of c-MYC, and we observed an increased incident of cytosolic *S.* Typhimurium in 10074-G5-treated cells, aligning with that observed in siRNA experiment (Figure 2C).

**Figure 2.**
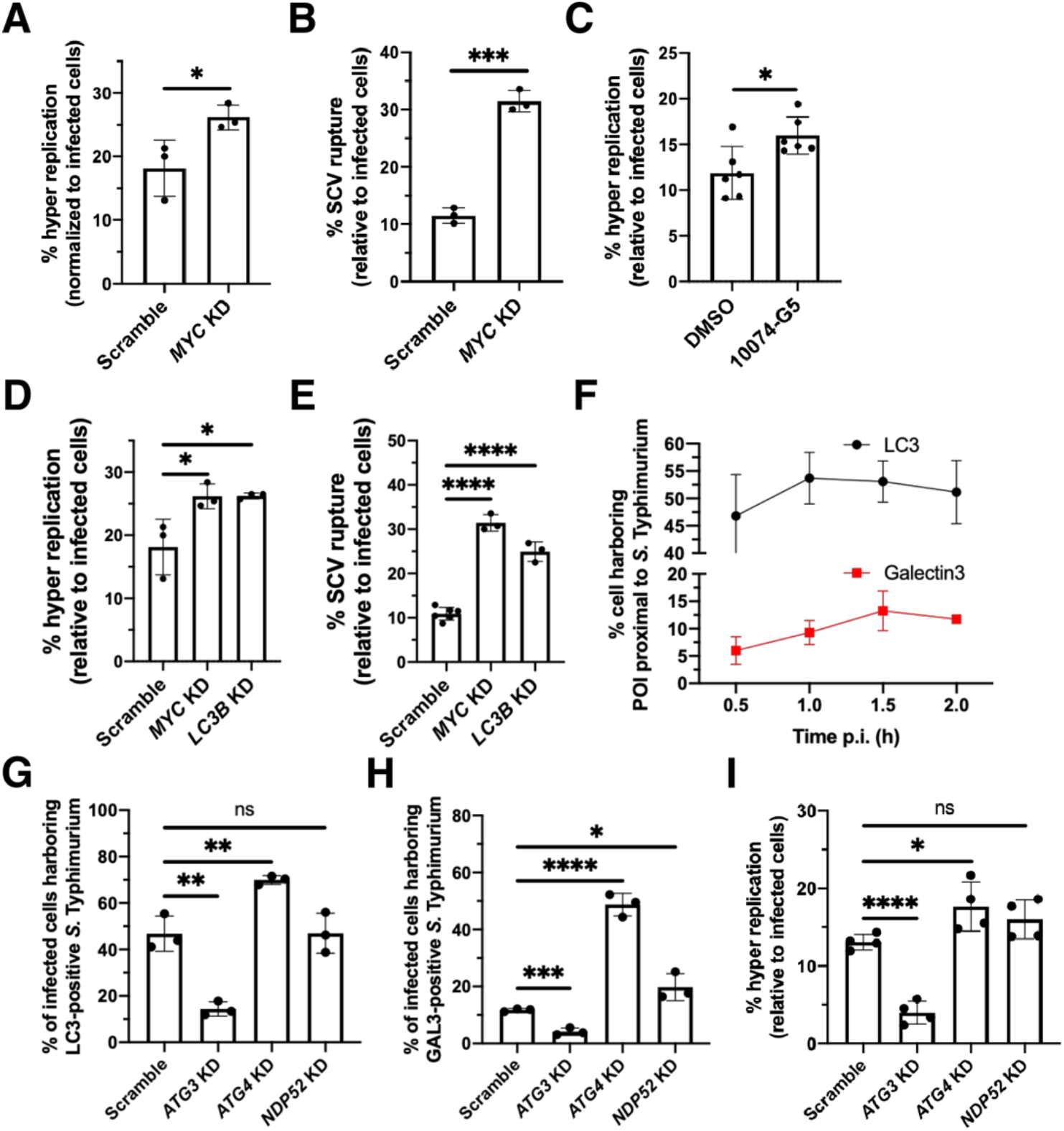
Host c-MYC and LC3 suppress *S.* Typhimurium cytosolic lifestyle in epithelial cells. (A) Quantification of the incidence of *S.* Typhimurium hyper replication under *c-MYC* knockdown at 6h pi by flow cytometry (n = 3). (B) Quantification of the incidence of SCV rupture under *c-MYC* knockdown at 2h pi by immunofluorescence staining of Galectin 3 (n = 3). (C) Quantification of the incidence of *S.* Typhimurium hyper replication under c-MYC inhibition at 6h pi by flow cytometry (n = 5). (D) Quantification of the incidence of *S.* Typhimurium hyper replication under *c-MYC* and *LC3B* knockdown at 6h pi by flow cytometry (n = 3). (E) Quantification of the incidence of SCV rupture under *c-MYC* and *LC3B* knockdown at 2h pi by immunofluorescence staining of Galectin 3 (n = 3). (F) Quantification of LC3 and Galectin 3 proximal to *S*. Typhimurium at 0.5h, 1h, 1.5h and 2h pi by immunofluorescence staining (n = 3). (G) Quantification of LC3 proximal to *S*. Typhimurium under *ATG3*, *ATG4* and *NDP52* knockdown at 0.5h pi by immunofluorescence staining (n = 3). (H) Quantification of Galectin 3 proximal to *S*. Typhimurium under *ATG3*, *ATG4* and *NDP52* knockdown at 2h pi by immunofluorescence staining (n = 3) (I) Quantification of *S.* Typhimurium hyper replication under *ATG3*, *ATG4* and *NDP52* knockdown at 6h pi by flow cytometry (n = 3). Significance cut-offs: ns = not significant/ *P* ≥ 0.05, * = *P* < 0.05, ** = *P* < 0.01, *** = *P* < 0.001 and **** = *P* < 0.0001.

### c-MYC restricts *S.* Typhimurium cytosolic access via downstream-regulated LC3

Given the established roles of c-MYC as transcription as well as cell cycle regulator, we aimed to determine whether the c-MYC downstream regulator of *S*. Typhimurium lifestyle is part of either the c-MYC regulon, or rather a cell cycle-dependent gene that is indirectly regulated by c-MYC. To test these two possibilities, we synchronized the cell using double-thymidine block and release at designated time prior to *S.* Typhimurium infection (Figures S5A – S5C). With cells synchronized to G1, G2/M and S phase, we did not observe a significant change in the incidence of cytosolic *S.* Typhimurium, suggesting c-MYC regulates *S.* Typhimurium cytosolic access independent of the host cell cycle (Figure S5D).

To decipher the underlying regulatory mechanism of c-MYC on *S.* Typhimurium cytosolic lifestyle, we looked into the c-MYC regulon. Among the c-MYC regulated genes, we identified microtubule-associated protein 1A/1B-light chain 3 (LC3) that has been closely associated with restriction of cytosolic bacteria (Siqueira et al., 2018). It was reported that c-MYC knockdown in HeLa cells leads to a reduced LC3 level as well as autophagosome and lysosome formation (Annunziata et al., 2019; Toh et al., 2013). The selection of LC3 by a candidate approach was also coherent to the range of autophagic genes detected among our screen candidates in our study, as well as the established role of autophagy on *S*. Typhimurium lifestyle. To determine whether LC3 is one of the c-MYC-regulated genes that limits *S.* Typhimurium cytosolic lifestyle, we performed siRNA knockdown of *LC3B* and assayed the incidence of cytosolic *S.* Typhimurium and SCV rupture. Under *LC3B* knockdown, we observed an increased level cytosolic *S.* Typhimurium and SCV rupture aligning with that of c-MYC knockdown (Figures 2D and 2E). Beside knocking down c-MYC, we also tested the impact of GFP-LC3B over-expression on *S.* Typhimurium lifestyles. Under GFP-LC3B over-expression, an elevated level of cytosolic *S.* Typhimurium was observed, indicating LC3B level is a significant factor contributing to *S.* Typhimurium cytosolic lifestyle (Figures S6 and S7). Together, the identified roles of c-MYC and LC3 in restricting cytosolic *S.* Typhimurium also align with the detection of multiple xenophagy genes among the screen hits that were filtered out from our screening results due to the stringency applied (data not shown). Despite demonstrating LC3 serves as a c-MYC downstream regulated gene that modulates cytosolic *S.* Typhimurium, this observation suggests another layer of regulation at SCV stability, in addition to the currently perceived role of LC3 in restricting cytosolic bacteria via autophagy.

### LC3 is recruited to SCV prior to catastrophic damage independent of NDP52

To resolve how LC3 contributes to regulate SCV stability, we first monitored the subcellular distribution of LC3 during early infection as elevated SCV rupture was reported at 2 h pi as indicated by Galectin 3. Previously reported pathways that regulates SCV stability in enterocytes also function in early infection, suggesting the fate of SCV is determined within early hours of infection (Fredlund et al., 2018; Stévenin et al., 2019). By simultaneously tracking the distribution of Galectin 3 and LC3 in the host cell during the first two hours of infection, we observed different kinetics of recruitment to SCV proximity. We observed an increase from 6% (30 min pi) to 13.2% (90 min pi) of Galectin 3 recruitment, whereas a stable frequency of LC3 between 45% to 55% throughout the first two hours (Figure 2F). The higher level of LC3 recruitment than Galectin 3 suggested that LC3 is being recruited to SCV before and independent of the exposure of host glycans that marks catastrophic endomembrane damage (Ellison et al., 2020).

A recent report has demonstrated the SCV becomes leaky during early infection, and these leaky SCV are LC3-positive (Xu et al., 2019). Calcium-binding and coiled-coil domain-containing protein 2 (NDP52), an adaptor protein that targets Galectin 8-bound damaged endomembranes to autophagosome by binding to Galectin 8 and LC3 (Thurston et al., 2012). To determine whether the recruitment of LC3 to SCV during early infection is dependent on NDP52, we performed siRNA knockdown of NDP52 and measured the LC3 recruitment, SCV rupture and cytosolic *S.* Typhimurium. Under NDP52 knockdown, we observed a slight increase in SCV rupture at 2 h pi, but indifferent levels of LC3 recruitment at 30 min pi and cytosolic *S.* Typhimurium at 6 h pi (Figures 2G - 2I). This argues that NDP52 is dispensable for the recruitment of LC3 to SCV proximity during early infection.

### LC3-PE conjugation promotes catastrophic SCV damage

As LC3 is recruited to leaky SCV during early infection in the absence of catastrophic membrane damage, the cause-effect relation between LC3 recruitment and SCV integrity remains elusive. In the biogenesis of autophagosome, Pro-LC3 was first cleaved by ATG4 to give LC3-I, which is sequentially conjugated with ATG7, ATG3 and ATG16L1 complex to ultimately conjugate with membrane-anchored phosphatidylethanolamine (PE) to form LC3-II (Randall-Demllo et al., 2013). LC3-II is then turned over by ATG4 to give LC3-I and PE (Agrotis et al., 2019). To determine whether the conjugation of LC3 to endomembrane is a regulatory factor of SCV integrity and cytosolic *S.* Typhimurium lifestyle, we blocked the LC3 conjugation and deconjugation by knocking down ATG3 and ATG4, respectively. We first confirmed the silencing of ATG3 and ATG4 will respectively decrease and increase the LC3 recruitment towards SCV proximity (Figures 2G). We then measured the frequency of SCV catastrophic damage and cytosolic *S.* Typhimurium under ATG3 and ATG4 knockdown. When ATG3 is silenced, we observed a significantly lower level of SCV rupture and cytosolic *S.* Typhimurium, while ATG4 knockdown yielded higher levels in both readouts (Figures 2H and 2I). The opposite phenotypes led by the increment and reduction of LC3-PE conjugation indicates the membrane conjugation of LC3 act as a promoting factor for SCV catastrophic damage and subsequent cytosolic *S.* Typhimurium lifestyle.

### *S.* Typhimurium SopF avoids LC3 recruitment to evade catastrophic SCV damage and modulate host inflammation level by balancing intracellular lifestyle

With the recognition of LC3-PE conjugation being a promoting factor of SCV catastrophic damage, and a previously reported *S.* Typhimurium T3SS1 effector, SopF that blocks LC3 conjugation on SCV and limits SCV damage (Lau et al., 2019; Xu et al., 2019). We hypothesized that the *S.* Typhimurium employs SopF to inhibit LC3 conjugation on SCV membrane as a strategy to modulate the level of cytosolic *S.* Typhimurium. To test this hypothesis, we first confirmed the role of SopF in limiting LC3 recruitment towards SCV prior to SCV rupture by fluorescent microscopy. At 30 min pi, we observed an elevated recruitment of LC3 but not Galectin 3 in Δ*sopF* mutant, suggesting SopF inhibits LC3 recruitment prior to SCV rupture, at the same time confirming the observation by Lau et al. 2019 (Figures 3A – 3C). To determine whether SopF acts to limit SCV rupture by inhibiting LC3 recruitment, we used the Δ*sopF* mutant in conjunction with LC3 lipidation inhibition by ATG3 knockdown. We observed reduced levels of LC3 recruitment, SCV rupture and cytosolic *S.* Typhimurium lifestyle of the Δ*sopF* mutant under ATG3 KD, suggesting the Δ*sopF*-induced SCV rupture requires LC3 lipidation (Figures 3D – 3F).

**Figure 3.**
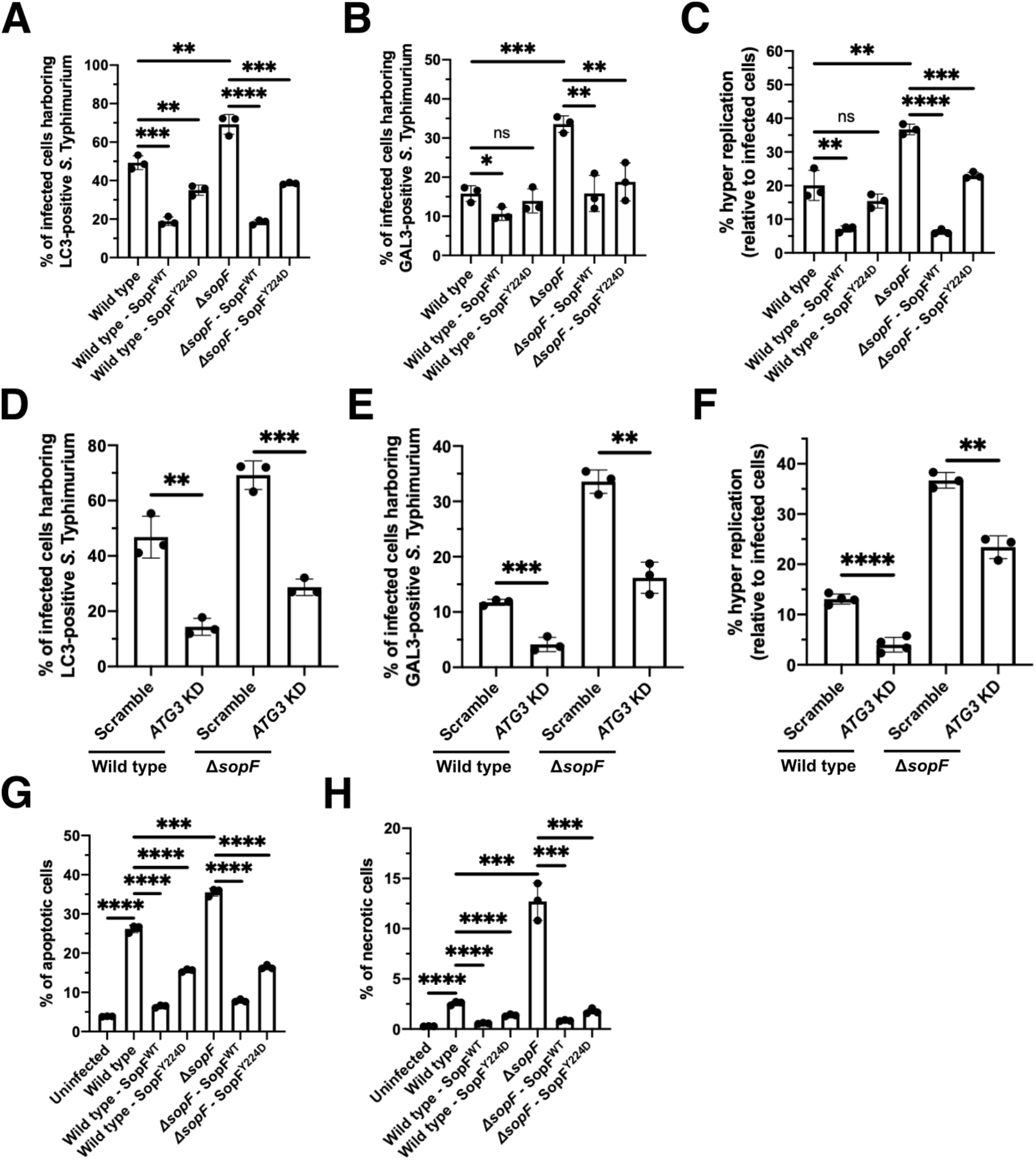
*S.* Typhimurium SopF regulates *S.* Typhimurium cytosolic lifestyle via SCV stabilization in a dosage-dependent manner. (A) Quantification of LC3 proximal to different *S*. Typhimurium stains at 0.5h pi by immunofluorescence staining (n = 3). (B) Quantification of Galectin 3 proximal to different *S*. Typhimurium strains at 2h pi by immunofluorescence staining (n = 3) (C) Quantification of *S.* Typhimurium hyper replication in different *S*. Typhimurium stains at 6h pi by flow cytometry (n = 3). (D) Quantification of LC3 proximal to wild type and Δ*sopF S*. Typhimurium under *ATG3* knockdown at 0.5h pi by immunofluorescence staining (n = 3). (E) Quantification of Galectin 3 proximal to wild type and Δ*sopF S*. Typhimurium under *ATG3* knockdown at 2h pi by immunofluorescence staining (n = 3) (F) Quantification of wild type and Δ*sopF S.* Typhimurium hyper replication under *ATG3* knockdown at 6h pi by flow cytometry (n = 3). (G) Quantification of the apoptotic cell abundance upon infection by different *S*. Typhimurium stains at 6h pi by Caspase 3/7 activity staining and flow cytometry (n = 3). (H) Quantification of the necrotic cell abundance upon infection by different *S*. Typhimurium stains at 6h pi by SYTOX AADvanced staining and flow cytometry (n = 3).Significance cut-offs: ns = not significant/ *P* ≥ 0.05, * = *P* < 0.05, ** = *P* < 0.01, *** = *P* < 0.001 and **** = *P* < 0.0001.

To address whether *S.* Typhimurium employs SopF to regulate SCV stability in a dosage dependent manner, we discretely overexpressed SopF in *S.* Typhimurium with SINA variants and ectopically overexpressed eGFP-SopF in HeLa cells, and assayed the LC3 recruitment, SCV damage and cytosolic *S.* Typhimurium (Figure S8). From the parallel experiments, we observed a reciprocal effect of SopF over-expression and *sopF* knockout, where high levels of SopF led to reduced cytosolic *S.* Typhimurium and vice versa (Figures 4A – 4C and S9). The dosage-dependent effects of SopF on LC3 recruitment, SCV integrity and cytosolic *S.* Typhimurium argue that SopF secretion dosage is a plausible strategy of *S.* Typhimurium in modulating the balance of intracellular lifestyle.

**Figure 4.**
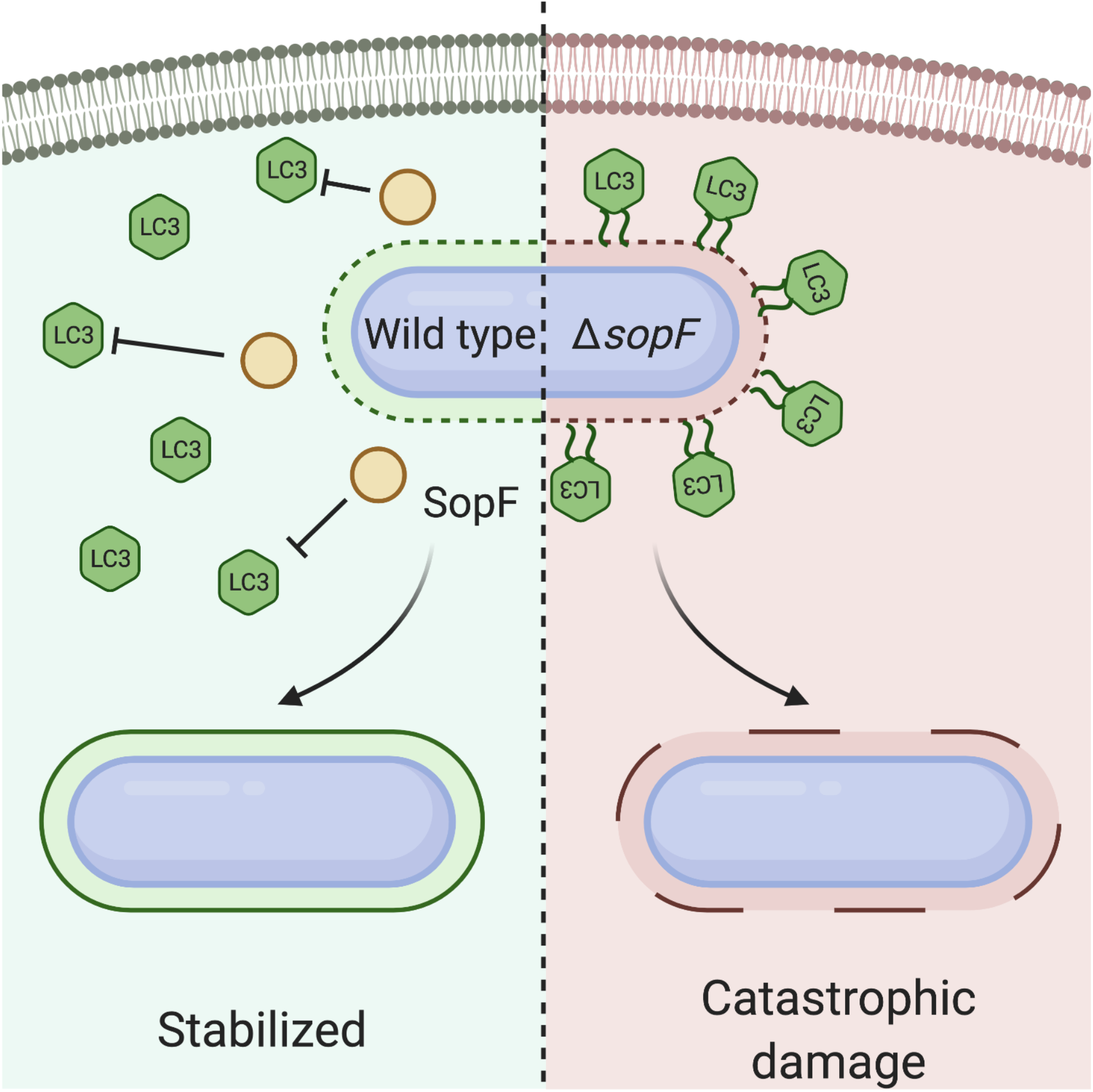
Cellular model of *S.* Typhimurium subversion strategy to avoid targeted by host autophagy. Shortly after bacterial uptake by the host, SCV turns porous and is targeted by host autophagy machinery. The lipidation and delipidation of autophagy protein LC3 control the LC3 association with SCV, which promote the catastrophic damage of SCV and expose *S.* Typhimurium to host cytosolic autophagy. *S.* Typhimurium employs T3SS1 effector SopF to impede the LC3 recruitment towards SCV as a strategy to balance the incidence of different *S.* Typhimurium intracellular lifestyles and the consequent host cell fate.

To decipher the physiological implication of *S.* Typhimurium SopF-mediated LC3 subversion and lifestyle balancing, we assayed the infection-induced host cell death by measurement of cell membrane integrity and activity of apoptotic Caspases. We observed an elevated level of lytic cell death and apoptosis in Δ*sopF* mutant as compared wild type, while reduced lytic cell death and apoptosis were observed in SopF^WT^ and SopF^Y224D^ over-expressing conditions (Figures 3G and 3H). Such observations were also coherent with the increased abundance of deformed dead cells detected (Figure S10). Altogether, our observations suggested the role of *S.* Typhimurium SopF in modulating LC3 recruitment to SCV as a strategy to balance intracellular lifestyle and host cell death (Figure 4).

## Discussion

*S.* Typhimurium has been reported to undergo different adaptation on its metabolism as well as virulence marker expression to get through the highly heterogeneous microenvironments of the host and cause a range of pathological consequences. Several host endocytic trafficking machineries have been identified to determine the intracellular fate of *S.* Typhimurium, yet alternative contributors from other host pathways remain unidentified. Combining the SINA reporter system and the Gene Trap forward genetic screen, we identified the oncogenic transcription factor, c-MYC as a negative regulator of cytosolic *S.* Typhimurium lifestyle. Herein, we reported that LC3, a downstream regulated gene of the c-MYC cascade, mediates the restriction function on *S.* Typhimurium cytosolic lifestyle. We further demonstrated the functional significance of LC3 lipidation on SCV catastrophic damage and a dose-dependent modulation of *S.* Typhimurium lifestyle by T3SS1 effector SopF via subverting LC3 recruitment to SCV.

In this work, we employed the Gene Trap screening platform to identify the host factors that restrict cytosolic *S.* Typhimurium lifestyles. The setup of the screen offers a number of merits as compared to classical image-based screening approaches, including possibility to target non-coding regions, additional enrichment and reduced demand on labor. The Gene Trap adopts a retroviral vector that randomly integrates at transcription start site proximity of active genes (Ambrosi et al., 2011). The retrovirus integration not only allows us to target coding genes, but also non-coding sequences that were also targeted in our screen. This allows the exploration of non-coding that is less readily achieved by most reverse genetic screens. Due to the low permissiveness of cytosolic *S.* Typhimurium in eHAP cells, increment in cytosolic *S.* Typhimurium can be enriched with greater confidence, which enabled us to obtain hits with fold-change in the range of hundreds that are verified in independent experiments (data not shown). The Gene Trap approach is coupled with flow cytometry, which reduces the labor-intensive pipetting and imaging process that are critical for imaged-based screening, ultimately enabling a higher throughput workflow. Among the candidate genes in enriched samples, we detected a number of xenophagy-related candidates, including the LC3-II delipidating enzyme ATG4C, further supporting the functionality of our screen in identifying host factors that limit cytosolic *S.* Typhimurium. We also identified candidates from the Syntaxin family, which has been implied in SCV stability in our previous work, offering additional confidence on the array of approaches used to probe *S.* Typhimurium lifestyles (Stévenin et al., 2019). Due to our highly stringent filtering parameters and limited sequencing depth, some of the above-described candidates did not show up in the filtered list.

The identification of c-MYC as a regulator of *S.* Typhimurium lifestyle enriches the understanding on the regulatory mechanism on *S.* Typhimurium subcellular localization in enterocytes. Previous work our team has revealed the intracellular trafficking pathways and machinery that contributes to SCV integrity during early infection (Fredlund et al., 2018; Kreibich et al., 2015; Santos and Enninga, 2016; Stévenin et al., 2019). In the context of in vivo infections, there are a number of factors that are not completely recapitulated in in vitro cell culture systems, including heterogeneity in cellular stemness, three-dimension organization and mechanical traits. With our findings, we drew a potential link between cellular transcription profile and the intracellular bacterial physiology. At the native site of enteric bacteria, the intestine offers a highly-folded tissue organization comprised of intestinal epithelial stem cells at the crypts and enterocytes that differentiate and pushed towards the tip of the villus (Peterson and Artis, 2014). These intestinal epithelial cells harbor different lineage commitment and expression profile, which could also be of distinct preference for intracellular pathogen. Bacterial preference has been proposed, yet a discussion of bacterial preference in the context of *S.* Typhimurium lifestyles remains needed, where intracellular environment has been regarded as a crucial determinant factor for the life and death of pathogens.

Autophagy has been well-documented to target and eradicate the cytosolic bacteria via targeting the disrupted cellular membrane structures and ubiquitinated bacteria, serving as one of the host cellular machineries that regulates *S.* Typhimurium lifestyle (Wu et al., 2020). Autophagy has been implied to restrict cytosolic *S.* Typhimurium, while independent reports have offered opposite insight on the role of autophagy for *S.* Typhimurium cytosolic lifestyle. Such discrepancy could be explained by the different approaches applied to inhibit autophagy in the host cell as well as the experimental readouts selected (Birmingham et al., 2006; Yu et al., 2014). In our study, we employed multiple functional reporter readouts to directly depict the SCV rupture and cytosolic *S.* Typhimurium at corresponding timepoints. Our observation on LC3 recruitment prior to SCV rupture put forward the evidence of LC3 recruitment to the SCV proximity independent of SCV rupture and autophagophore adaptor NDP52. This suggests an alternative machinery that promoted the recruitment of LC3 in the early stage of *S.* Typhimurium infection prior to SCV rupture. We determined that the LC3-lipidating enzyme ATG3 and LC3-delipidating enzyme ATG4 balance the LC3 recruitment and respectively increase and decrease SCV rupture, in agreement with our detection of ATG4C as a candidate of restricting *S.* Typhimurium cytosolic lifestyle. Our deduction on the involvement of LC3 lipidation for SCV stability proposed an additional role of LC3 in restricting *S.* Typhimurium intracellular survival, where LC3 drives SCV rupture and releases *S.* Typhimurium into the host cytosol to be targeted by autophagy. This model agrees with the previous observation on the reduced intracellular survival of Δ*sopF* mutant, which fails to avoid LC3 recruitment towards SCV (Xu et al., 2019).

The subcellular localization of *S.* Typhimurium in enterocytes serves as significant cues for the host cellular immunity, where vacuolar *S.* Typhimurium has been regarded as a more subtle form that evades the host immunity, while cytosolic *S.* Typhimurium is described to activate NLRP3 inflammasome and trigger host pyroptosis (Knodler et al., 2010, 2014; Sellin et al., 2014). The fate of infected host cells holds a critical role on the successful containment of *S.* Typhimurium infection as pyroptotic cells release IL-1β, IL-18 as well as cellular contents that signify tissue inflammation and recruit neutrophils for bacterial clearance (Sahoo et al., 2011). To achieve colonization, *S.* Typhimurium has to balance its intracellular lifestyle to enable efficient colonization and spreading while avoiding excessive inflammation and recruitment of immune cells. Previous reports have suggested SCV becomes porous in early infection, which is subsequently sensed by the V-ATPase and targeted by autophagy (Xu et al., 2019). This autophagy targeting is impeded by *S.* Typhimurium SopF via ATP6V0C ADP-ribosylation, and we offered the evidence that SopF in turn inhibits the recruitment of the SCV disruptor, LC3, in a dosage-dependent manner to modulate *S.* Typhimurium intracellular lifestyle.

Altogether, we demonstrated the versatility of fluorescent reporters in host-pathogen interactions studies, the putative link between cellular status and bacterial pathogenesis, and an unprecedent role of autophagy in restricting cytosolic *S.* Typhimurium as well as the countermeasures of *S.* Typhimurium in mitigating successful survival in hosts.

## Acknowledgements

We thank the members of the Dynamics of Host-Pathogen Interactions Unit for the constructive comment and discussion. We are grateful for the technical support by Laurence MA, Christiane BOUCHIER and Cédric FUND from the BIOMICS platform, Institut Pasteur. This research was supported by fellowships from Croucher Foundation (HK) and Fondation pour la Recherche Médicale (FR) to C.H.L. C.H.L. is part of the Pasteur - Paris University (PPU) International PhD Program. J.E. is supported by the ERC-CoG “Endosubvert”. The Enninga lab is part of the LabEx IBEID and Milieu Interieur.

## Author Contributions

C.H.L. and J.E. conceptualized the project, C.H.L. conducted the investigation, W.Y. performed the informatic analysis, C.H.L. and J.E. prepared and reviewed the manuscript.

## Materials and Methods

### Mammalian cell culture

HeLa cervical adenocarcinoma cells and HEK293T embryonic kidney cells were purchased from American Type Culture Collection (ATCC) and used within 20 passages of receipt. eHAP engineered human chronic leukemia cells were purchased from Horizon Genomics and used within 10 passages upon receipt. HeLa and HEK293T cells were cultured in Dulbecco’s Modified Eagle Medium (DMEM, high glucose, GlutaMAX™ Supplement, ThermoFisher) containing 10% (v/v) heat-inactivated fetal bovine serum (FBS, Sigma) and incubated at 37 °C with 5% CO_2_ and 100% humidity. eHAP cells were cultured in Iscove’s Modified Dulbecco’s Medium (IMDM, ThermoFisher) containing 10% (v/v) FBS and incubated at 37 °C with 5% CO_2_ and 100% humidity. For flow cytometry analysis, HeLa cells were seeded in 12-well tissue culture-treated plates (Corning Costar®) at a density of 9×10^4^ cells/well 72h prior to infection. For immunofluorescence staining, HeLa cells were seeded on UV-treated glass coverslips (Marienfeld) in 24-well plates at a density of 4×10^4^ cells/well 72h prior to infection. For infection of mutagenized library, mutagenized eHAP cells were seeded on 10 cm 10 cm tissue culture-treated dishes (Corning Costar®) at a density of 1.8×10^6^ cells/dish 48h prior to infection.

### Bacterial strains

Bacterial strains and plasmids used in this study are listed in Tables S3 and S4, respectively. All mutants were constructed using bacteriophage λ red recombinase system from parental stain *Salmonella* Typhimurium strain SL1344 using primers listed in Table S5 (Santiviago et al., 2009). Bacteria were cultured in Lysogeny broth (LB) supplemented with appropriate antibiotics, where necessary (Ampicillin 100 μg/mL; Kanamycin 50 μg/mL).

### Plasmid construction

The construction of pSINA1.1 and pSINA1.7 were described in our previous work (Luk et. al. 2020). *sopF* coding sequence was first amplified with sopF_fw and sopF_rv and replaced the smURFP-HO-1 sequence of pBAD smURFP-HO-1 to yield pBAD *sopF*^WT^. pBAD *sopF*^WT^ was used as the template for site-directed mutagenesis using primers Y224D_fw and Y224D_rv to introduce Y224D point mutation and generate pBAD *sopF*^Y224D^. The arabinose inducible cassette of *sopF*^WT^ and *sopF*^Y224D^ were amplified using ara_fw and ara_rv and inserted at the Nru I site of pSINA1.1 to generate pSIN1.1 PBAD *sopF*^WT^ and pSINA1.1 PBAD *sopF*Y224D.

To generate peGFP-*sopF*^WT^ and peGFP-*sopF*^Y224D^, *sopF*^WT^ and *sopF*^Y224D^ were respectively amplified from pBAD *sopF*^WT^ and pBAD *sopF*^Y224D^ using eGFP-sopF_fw and eGFP-sopF_rv, and inserted between EcoR I and Sal I sites of peGFP-C3.

### Haploid genetic screen for *S.* Typhimurium lifestyle regulators

Gene-trap retrovirus was produced in HKE293T cells using the four plasmids described by Jae et al., 2013. To produce gene-trap retrovirus, HEK293T cells were seeded on 20 10 cm dish at 60% confluent. In the following day, each plate of HEK293T cells was transfected with VSV-G (2.5 μg/plate), pAdvantage (10 μg/plate), gag-pol (10 μg/plate) and pGeneTrap (10 μg/plate) using CaCl_2_ transfection method (250 mM CaCl_2_, 1 x HEPES, pH 7.04). Cell medium was collected on day 2 and 3 post transfection, first centrifuged to remove cells, and subsequently ultra-centrifuged at 20000 g at 4°C for 2 hours. The viral particles were stored at −80 °C.

To generate the mutagenized eHAP library, 30 million eHAP cells were seeded and transduced with gene-trap retrovirus harvested from three combined harvest in the presence of 8 μg/mL protamine sulfate (Sigma). The transduced cells were expanded and passed for a period of 10 days upon transduction. The mutagenized libraries were maintained for at least 3-fold complexity.

The mutagenized cells were seeded and infected with *S.* Typhimurium harboring pSINA1.7, infected cells were harvested and subjected to cell sorting. After the enrichment of mutagenized eHAP harboring cytosolic *S.* Typhimurium, the cells were incubated in Qiagen buffer AL supplemented with Proteinase K at 56°C overnight to de-crosslink the cell pellet. In the following day, the genomic DNA were extracted from the cells using Qiagen DNA mini kit (Qiagen, USA) according to manufacturer’s manual and quantified using Nanodrop2000 spectrophotometer (ThermoFisher).

### Insertion site mapping

To map the insertion sites, genomic sequence flanking the insertion cassette were amplified using non-restriction linear amplification-mediated PCR (nrLAM-PCR) with a number of adaptation to GeneTrap approach (Paruzynski et al., 2010). For each nrLAM-PCR reaction, 1 μg of extracted genomic DNA was used as template for single-strand amplification. For each reaction, genomic DNA was combined with 0.01 mM dNTPs, 0.085 pmol double-biotinylated primer LTRI, Taq polyermase (1.25 U/rxn) and the supplied buffer (Qiagen). The reaction was performed for 50 cycles, with annealing at 58°C for 45 s and elongation at 72°C for 25s. Additional 2.5 U of Taq polymerase was supplemented to each reaction, and another 50 cycles were performed. To capture the biotinylated single-stranded DNA (ssDNA), each PCR reaction was combined in 1 to 1 ratio with DYNAL™ MyOne™ Dynabeads™ Streptavidin C1 (ThermoFisher) resuspended in 2 x binding buffer (6 M LiCl, 10 mM Tris, 1 mM EDTA, pH=7.5) for 2h at room temperature with horizontal shaking at 300 r.p.m. Prior to combination with PCR products, 20 μL of MyOne beads were washed once with 1 x PBS + 0.1% BSA and once with 1 x binding buffer (3 M LiCl, 5 mM Tris, 0.5 mM EDTA, pH=7.5). After capturing of ssDNA, the DNA-beads complex was resuspended in 100 μL of H_2_O.

For the ligation single-stranded linker, the DNA-beads complex was resuspended in a ligation mixture consisting of 10 pmol LC1, 25% PEG 8000, 1 mM hexamine cobalt (III) chloride, 1 mM ATP, 20 U T4 RNA ligase 1 and the supplied buffer (ThermoFisher), and incubated overnight at room temperature with horizontal shaking at 300 r.p.m. On the following day, the DNA-beads complex was washed with H2O (2X) and resuspended in 10 μL of H_2_O. Subsequent PCR amplification was performed with 5 μL of DNA-beads complex as template, combined with 0.2 mM dNTPs, 8.35 pmol biotinylated primer LTRII, 8.35 pmol LCI, Taq polyermase (1.25 U/rxn) and the supplied buffer (Qiagen). The reaction was performed for 35 cycles, with denaturation at 95°C for 45 s, annealing at 58°C for 45 s and elongation at 72°C for 25s. PCR products from each PCR reaction were captured using DYNAL™ MyOne™ Dynabeads™ Streptavidin C1. For each reaction, the non-biotinylated ssDNA was separated from the biotinylated ssDNA bound to magnetic beads by denaturation using fresh 0.1 N NaOH at room temperature for 10 minutes with horizontal shaking at 300 r.p.m. The second PCR amplification was performed with 5 μL of denatured non-biotinylated ssDNA as template, combined with 0.2 mM dNTPs, 8.35 pmol biotinylated primer LTRIII, 8.35 pmol LCII, Taq polyermase (1.25 U/rxn) and the supplied buffer (Qiagen). The reaction was performed for 35 cycles, with denaturation at 95°C for 45 s, annealing at 58°C for 45 s and elongation at 72°C for 25s. The PCR products were purified using AMPure XP beads according to manufacturer’s manual, and further ligated with illumina adaptor sequences using NEBNext^®^ Ultra™ II DNA Library Prep Kit for Illumina^®^ (New England Biolabs). The prepared libraries were purified using AMPure XP beads and sequenced as 2 × 250 bp on MiSeq Reagent Kit v2 (illumina) using sequencing primer Rd1 Seq Primer and Rd2 Seq Primer from illumina.

### Sequencing data processing

Paired-end sequencing reads with 250 nucleotides of each end in FASTQ files were decoded and adaptor sequence trimmed using by ea-utils tool (https://expressionanalysis.github.io/ea-utils/). Paired-end reads were merged into single end read using FLASH with at least 10 nucleotide overlap (Magoč and Salzberg, 2011). Merged reads were mapped to the human hg38 reference genome and genetrap sequence using BWA mem (with default settings. Mapped reads with less than 30 nucleotides of genetrap sequence were filtered out (Li and Durbin, 2010; Li et al., 2009). Among mapped read, the part of reads sequence mapped to human reference were annotated by HOMER (Heinz et al., 2010). The annotated reads were counted and classified into genetic elements including exon, intron, 5’ UTR and 3’ UTR using in-house Perl script.

### Analysis of screen hits

The insertion sites with less than 50 reads were first removed, hits falling into the coding sequencing of human genes were selected and the relative abundance of hits (read counts of gene X/total read counts) were calculated. Hits detected in at least 2 out of the 3 repeats were selected, and those exhibiting at least a 2-fold increase in enriched group (relative to unenriched control) were considered as significant for validation using siRNA gene silencing.

### Gene silencing by siRNA

The sequences of siRNA used were listed in Table S6, the siRNAs were ordered from Eurogentec (Eurogentech). Cells were seeded and transfected using Lipofectamin RNAiMAX (Thermofisher, USA) according to manufacturer’s manual 72 h prior to infection. Cell medium was replaced with fresh medium 24 h after transfection. siRNA knockdown efficiency was measured by Western Blot or qRT-PCR (Figure S11).

### Double-thymidine block

HeLa cells were seeded 72h on 12 wells plates prior to infection as described above. At 42h before infection, the cell culture medium was supplemented with 2 mM thymidine for 18h, cells were then washed and incubated in DMEM + 10% FBS for 8h. At 16h before infection, the cell culture medium was supplemented with 2 mM thymidine for 16h. The cells were then released by replacing the medium with DMEM + 10% FBS, cells synchronized at G1, S and G2/M phase were obtained at 0h, 3h and 6h post-release, respectively.

### Plasmid transfection

Plasmids used for transfection were listed in Table S4. Cells were seeded and transfected using XtrememGene 9 (Roche, USA) according to manufacturer’s manual 48 h prior to infection. Cell medium was replaced with fresh medium 24 h after transfection.

### Bacterial infections

Bacteria strains were streaked from glycerol stock on LB agar plates with appropriate antibiotics 2 days prior to infection. Three bacterial colonies were picked for overnight culture in LB medium supplemented with 0.3M NaCl with shaking at 37°C. 150 μL overnight culture was subculture in 3 mL LB + 0.3M NaCl (1:20 dilution) with shaking at 37°C for 3h. For strains harboring pSINA1.9, 0.1% L-arabinose was supplemented to the subculture 1 h before harvest. Bacteria were harvested with centrifugation (1 mL, 6000 x g, 1 min, RT), washed once in 1 x PBS and resuspended in DMEM with no FBS. HeLa cells were infected at a MOI of ~100 for 25 min at 37°C. Extracellular bacteria were removed and washed with 1 x PBS (5X). Cells were then incubated in DMEM + 10% FBS for 1h, washed with 1 x PBS (3X), incubated in DMEM + 10% FBS for 2 h, washed with 1 x PBS (3X) and then incubated in DMEM + 10% FBS supplemented with 10 μg/mL gentamicin for the remainder of the infection.

### Flow cytometry

At designated time points, cells were washed with 1 x PBS (1X) and detached with 0.05% Trypsin for 5 min at 37°C. Detached cells were mixed with equal volume of DMEM + 10% FBS, passed through 40 μm strainer and collected by centrifugation (500 x g, 5 min, 4 °C). Cell pellets were dislodged and fixed in 4% PFA (15 min, RT). Fixed cells were washed with 1 x PBS (2X) and resuspended in 200 μL 1 x PBS for further analysis. The fluorescent intensities of the samples were assayed with LSR Fortessa (BD) (tagBFP Ex: 405nm Em: 450/50nm; Timer^510^ Ex: 488nm Em: 525/50nm; Timer^580^ Ex: 562nm Em: 582/15nm; smURFP Ex: 633nm Em: 670/30nm) and analyzed with FlowJo (v10.0.4)

### Cell sorting

At 6 h pi, cells were washed with 1 x PBS (1X) and detached with 0.05% Trypsin for 5 min at 37°C. Detached cells were mixed with equal volume of DMEM + 10% FBS, passed through 40 μm strainer and collected by centrifugation (500 x g, 5 min, 4 °C). Cell pellets were dislodged and fixed in 4% PFA (15 min, RT). Fixed cells were washed with 1 x PBS (2X) and resuspended in 200 μL 1 x PBS for further analysis. Cells were sorted with Aria III (BD) (tagBFP Ex: 405nm Em: 450/50nm; Timer^510^ Ex: 488nm Em: 530/30nm; Timer^580^ Ex: 561nm Em: 586/15nm; smURFP Ex: 633nm Em: 660/20nm) to collect mutagenized eHAP cells harboring cytosolic *S.* Typhimurium.

### Immunofluorescence microscopy

Infected cells on coverslips were washed with 1 x PBS (1X) and fixed in 4% PFA (8 min, RT). After washing with 1 x PBS (3X), cells were permeabilized and blocked in 1 x PBS, 20% FBS, 0.25% saponin (30 min, RT). Coverslips were washed with 1 x PBS (3X) and incubated with anti-LC3 and anti-Galectin3 primary antibodies diluted in 1 x PBS, 1% BSA (60 min, RT), and then washed with 1 x PBS (3X) and incubated with Cy5-conjugated goat anti-rat secondary antibodies and Alexa Flour 488-conjugated goat anti-rabbit secondary antibodies diluted in 1 x PBS, 1% BSA (60 min, RT). Stained coverslips were then washed with 1 x PBS (3X) and mounted on SuperFrost Plus microscope sides (Thermo Scientific) with ProLong™ Gold Antifade Mountant without DAPI (Invitrogen). Samples were imaged with Perkin Elmer Ultraview confocal spinning disk microscope equipped with Volocity software and a 20X/1.3 N.A. air objective. Images were analyzed with FIJI (NIH) and figures were prepared using Inkscape Ver. 1.0.0.

### qRT-PCR

Cells were harvested on the same day of S. Typhimurium infection, the cells were collected by centrifugation at 500 g for 5 min. The total RNA was extracted using RNeasy Mini Kit (Qiagen) according to manufacturer’s manual. 5 μg of total RNA was used for reverse transcription using SuperScript III Reverse Transcriptase (Thermofisher) according to manufacturer’s manual. The reverse-transcribed cDNA was used as template for qRT-PCR using iTaq Universal SYBR Green Supermix (Bio-Rad) with primers described in Table S5. The Ct values obtained were used to calculate the expression of targeted genes relative to housekeeping gene (GAPDH).

### Western Blot

Cells were harvested on the same day of *S.* Typhimurium infection, the cells were collected by centrifugation at 500 g for 5 min. Cells were then lysed using 2x Laemmli Sample buffer (Bio-Rad) supplemented with 2-mercaptoethanol and heated at 95 °C for 5 min. The lysed samples were run on Bis-Tris NuPAGE protein gels (Invitrogen) and transferred to nitrocellulose membrane using Trans-Blot Turbo Transfer System (Bio-Rad). The membrane was first blocked in 1 x TBST (20 mM Tris-HCl, 150 mM NaCl and 0.1% (w/v) Tween 20), 2% BSA, and blotted with primary antibody (rabbit anti-c-MYC) in 1 x TBST, 1% BSA for 2 hours at room temperature. The membrane was then washed with 1 x TBST, 1% BSA (3X) for 15 min at room temperature. The membrane was then blotted with secondary antibody (goat anti-rabbit, HRP conjugate) for 2 hours at room temperature. The membrane was then washed with 1 x TBST, 1% BSA (3X) for 15 min at room temperature and incubated with ECL Western Blot Substrate (Thermofisher) and visualized using Azure c600 Gel Imaging System (Azure Biosystems). The membrane was then stripped using ReBlot Plus mild antibody stripping solution (Sigma), sequentially blotted with primary (mouse Anti-β-Actin) and secondary (goat anti-mouse, HRP conjugate) antibodies and visualized after ECL incubation.

### Caspase activity and cell death assay

At 6h pi, cells were detached using 0.05% trypsin and collected by centrifugation at 500 g for 5 min. The cells were then resuspended in DMEM + 10% FBS supplemented with 50 μM CellEvent™ Caspase 3/7 Green Detection Reagent (Invitrogen) at 37 °C for 25 min. 0.1 mM of SYTOX™ AADvanced™ Dead Cell Stain were then supplemented and incubated at at 37 °C for 5 min. The cells were immediately analyzed by flow cytometry. The live, apoptotic and necrotic cells were gated using the gating strategy described in Figure S10A.

### Statistical analysis

Unless further specified in the figure legend, data were analyzed for statistical significance with a Mann-Whitney test using Prism 8.0 (GraphPad). *P* value of < 0.05 is considered statistically significant. * = *P* < 0.05, ** = *P* < 0.01, *** = *P* < 0.001, **** = *P* < 0.0001, ns = not significant/ *P* ≥ 0.05.

---

**Figure S1.**
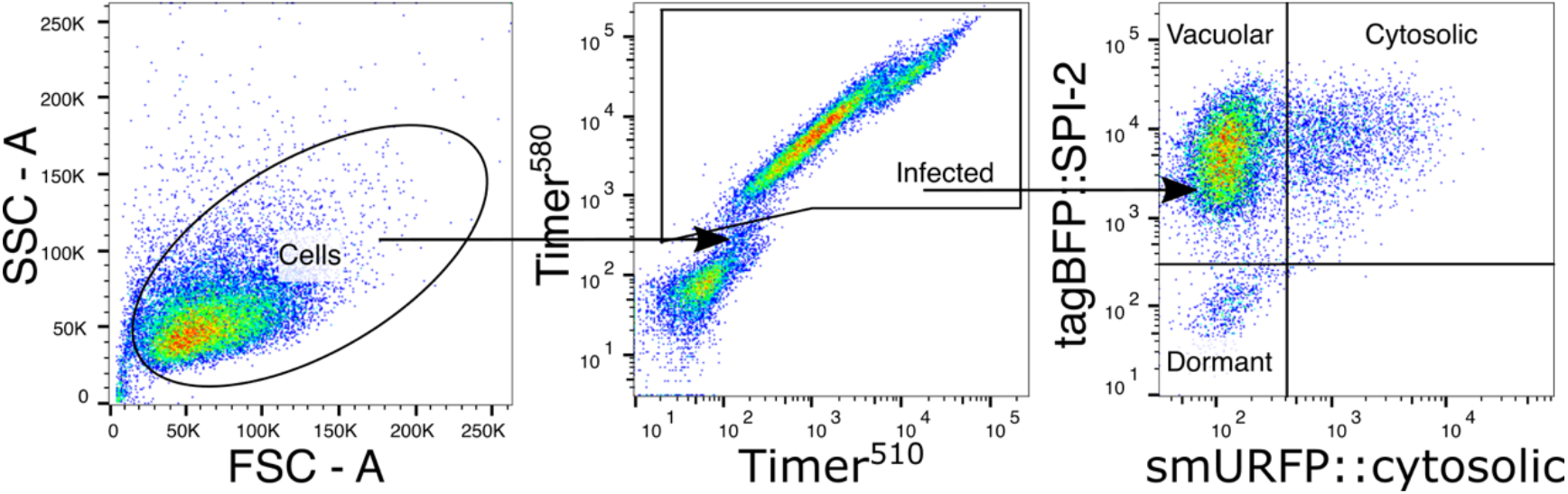
Gating strategy of SINA1.1 in HeLa cells. HeLa cells were infected with SINA1.1-haboring *S.* Typhimurium, harvested at 1h and 6h pi and analyzed by flow cytometry. The cells were first gated from the total recorded events on the SSC-A vs FSC-A plot to remove cell debris. The infected cells (infected) were gated by double-positive on Timer^580^ vs Timer^510^ plot. To determine the basal levels of P_*ssaG*_-tagBFP and P_*uhpT*_-smURFP, the sample harvested at 1h pi was used to define the basal levels and four quadrants were defined on the tagBFP::SPI-2 vs smURFP::cytosolic plot from “infected”.

**Figure S2.**
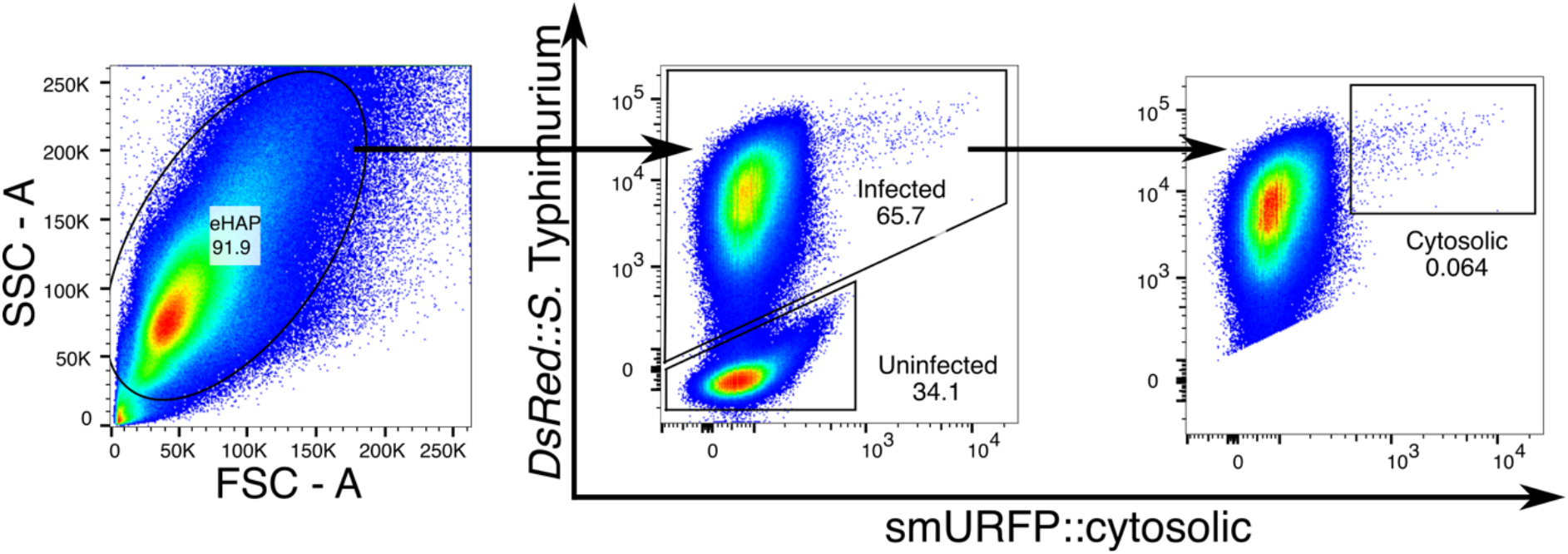
Gating strategy of SINA1.2 in eHAP cells. eHAP cells were infected with SINA1.2-harboring *S.* Typhimurium, harvested at 1h and 6h pi, analyzed by flow cytometry and sorted. The cells were first gated from the total recorded events on the SSC-A vs FSC-A plot to remove cell debris. The infected cells (infected) were gated by positive DsRed from P_*ybaJ*_-DsRed. The infected eHAP cells harboring cytosolic *S.* Typhimurium were gated by DsRed and smURFP double-positive on DsRed::*S.* Typhimurium vs smURFP::cytosolic plot.

**Figure S3.**
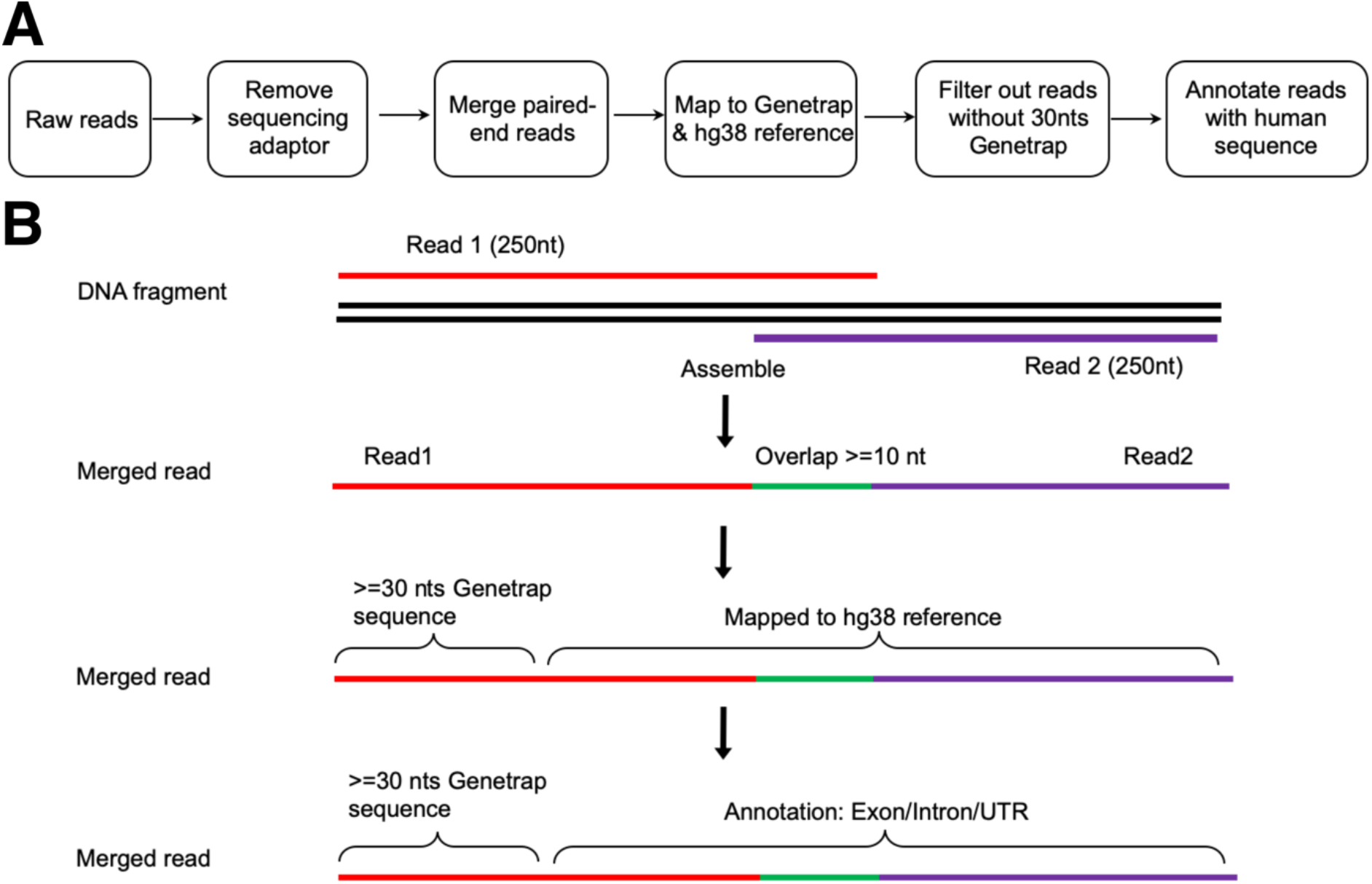
Illustration of sequencing analysis pipeline. (A) Workflow of NGS raw reads treatment and analysis. (B) Schematic illustration of the sequential steps for NGS raw reads treatment and analysis.

**Figure S4.**
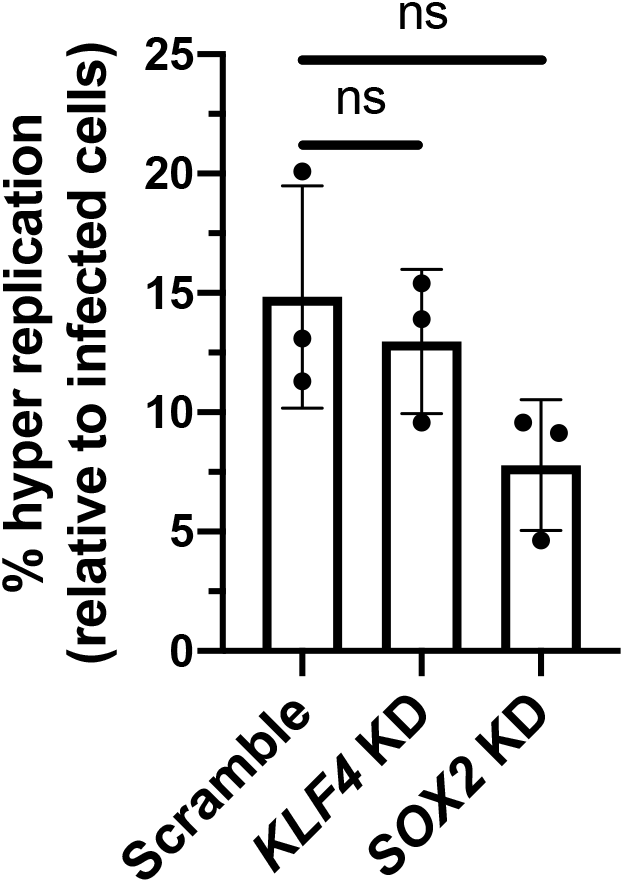
SOX2 and KLF4 are not regulators of cytosolic *S.* Typhimurium in HeLa cells. Quantification of the abundance of *S.* Typhimurium hyper replication in infected siRNA-transfeced HeLa cells at 6h pi. Significance cut-offs: ns = not significant/ *P* ≥ 0.05.

**Figure S5.**
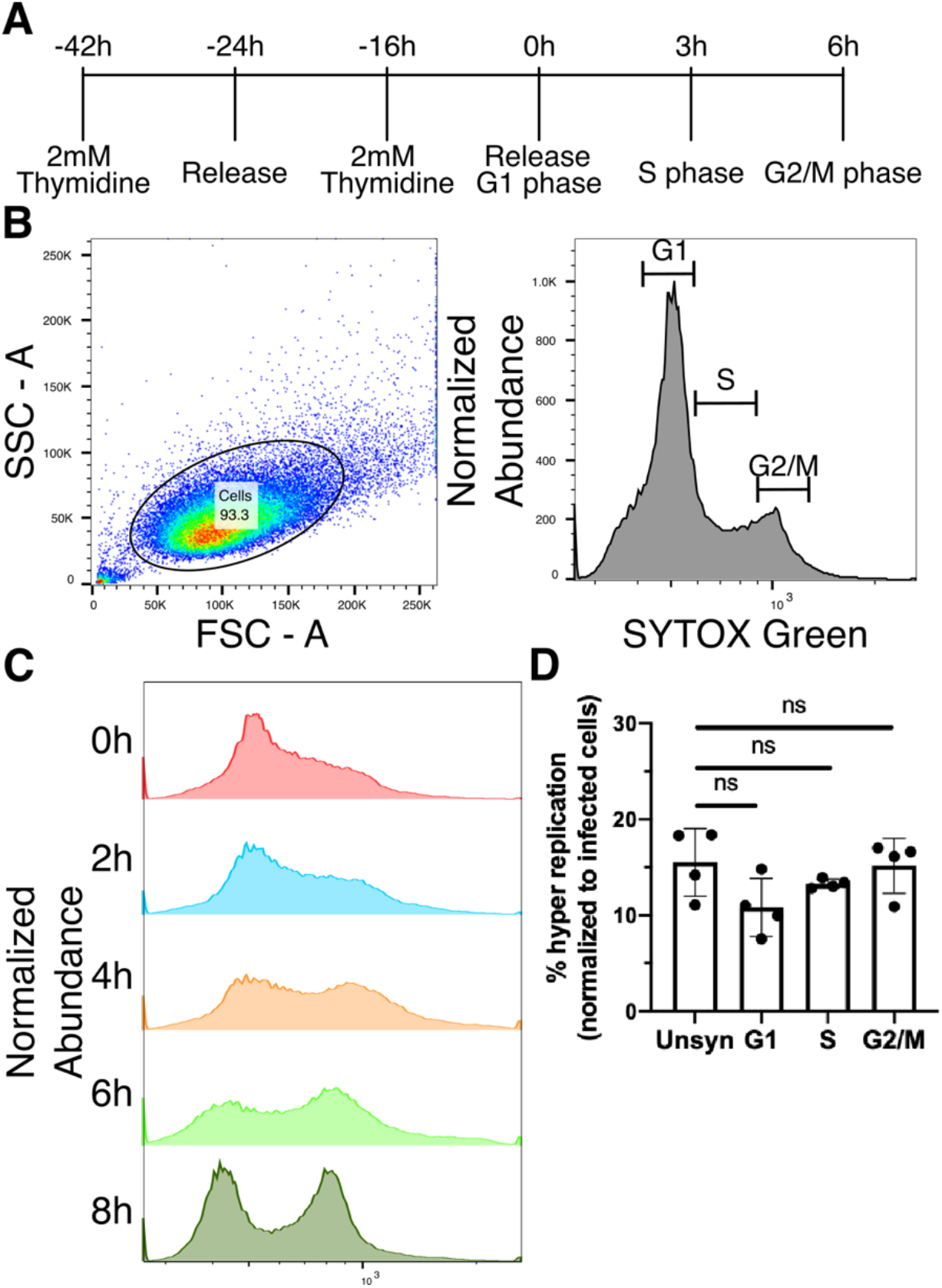
Cell cycle of HeLa cells is not a determinant factor for *S.* Typhimurium lifestyle. (A) Schematic illustration of cell cycle synchronization by double-thymidine block treatment. Illustration indicated the periods of thymidine blocks and the time required to reach each cell cycle upon release. (B) Gating strategy of cell cycle analysis. Cell were first gated from the total recorded events on the SSC-A vs FSC-A plot to remove cell debris, which were then displayed on histogram of SYTOX Green for cellular DNA content. (C) Cell cycle analysis of synchronized HeLa cells at 0h, 2h, 4h, 6h and 8h post-release. (D) Quantification of the abundance of *S.* Typhimurium hyper replication in infected synchronized HeLa cells at 6h pi. Significance cut-offs: ns = not significant/ *P* ≥ 0.05.

**Figure S6.**
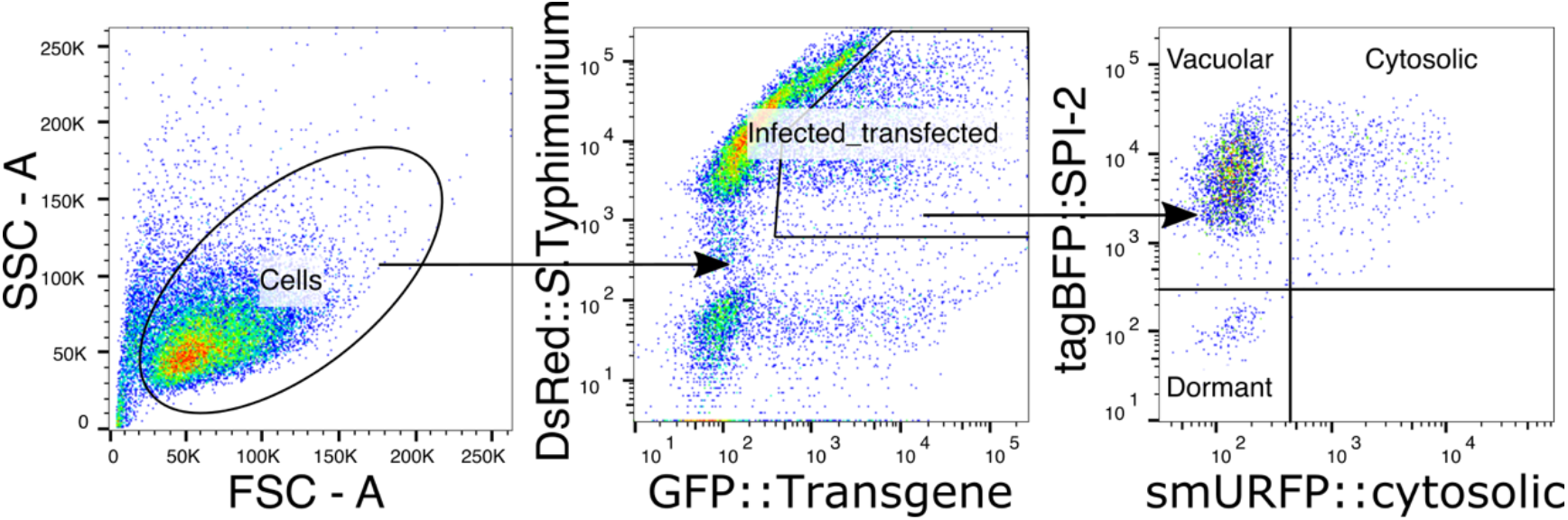
Gating strategy of SINA1.7 in combination with GFP-tagged transgene expression in HeLa cells. Transfected HeLa cells were infected with SINA1.7-haboring *S.* Typhimurium, harvested at 1h and 6h pi and analyzed by flow cytometry. The “cells” were first gated from the total recorded events on the SSC-A vs FSC-A plot to remove cell debris. The infected-transfected cells were gated by double-positive on DsRed::*S.* Typhimurium vs GFP::Transgene plot. To determine the basal levels of P_*ssaG*_-tagBFP and P_*uhpT*_-smURFP, the sample harvested at 1h pi was used to define the basal levels and four quadrants were defined on the tagBFP::SPI-2 vs smURFP::cytosolic plot from “infected-transfected”.

**Figure S7.**
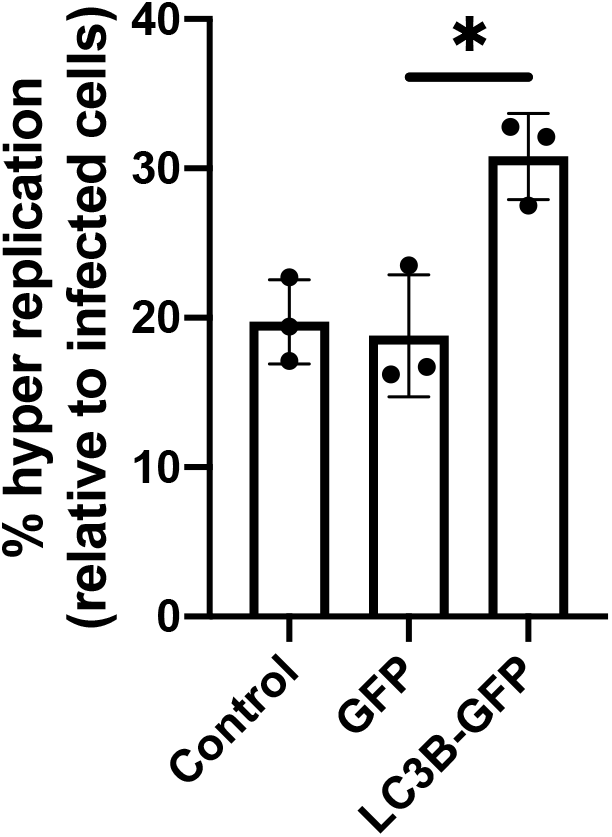
Ectopic LC3B expression elevates cytosolic *S.* Typhimurium in HeLa cells. Quantification of the abundance of *S.* Typhimurium hyper replication in transgene-expressing infected HeLa cells at 6h pi using flow cytometry (n = 3). Significance cut-offs: * = *P* < 0.05

**Figure S8.**
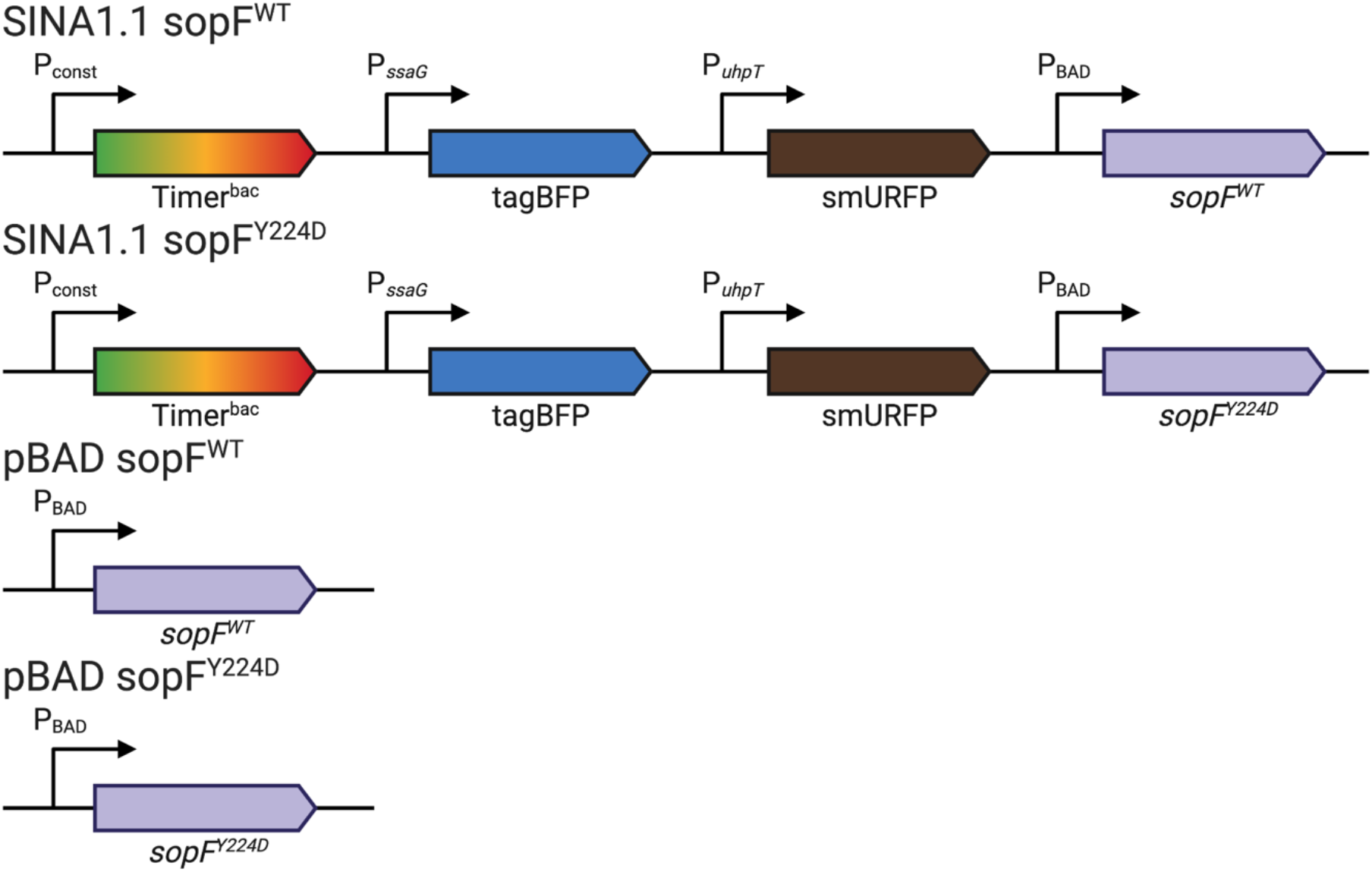
Schematic diagram of plasmid constructs used for Δ*sopF* complementation. Schematic illustration of plasmid constructs used for Δ*sopF* complementation and SopF over-expression. For the quantitative analysis of the distribution of *S.* Typhimurium intracellular lifestyle, SINA1.1 sopF^WT^ and SINA1.1 sopF^Y224D^-harboring *S.* Typhimurium were used in combination with flow cytometry. For immunofluorescent staining and Caspase activity and cell death measurement, pBAD sopF^WT^ and pBAD sopF^Y224D^-harboring *S.* Typhimurium were used.

**Figure S9.**
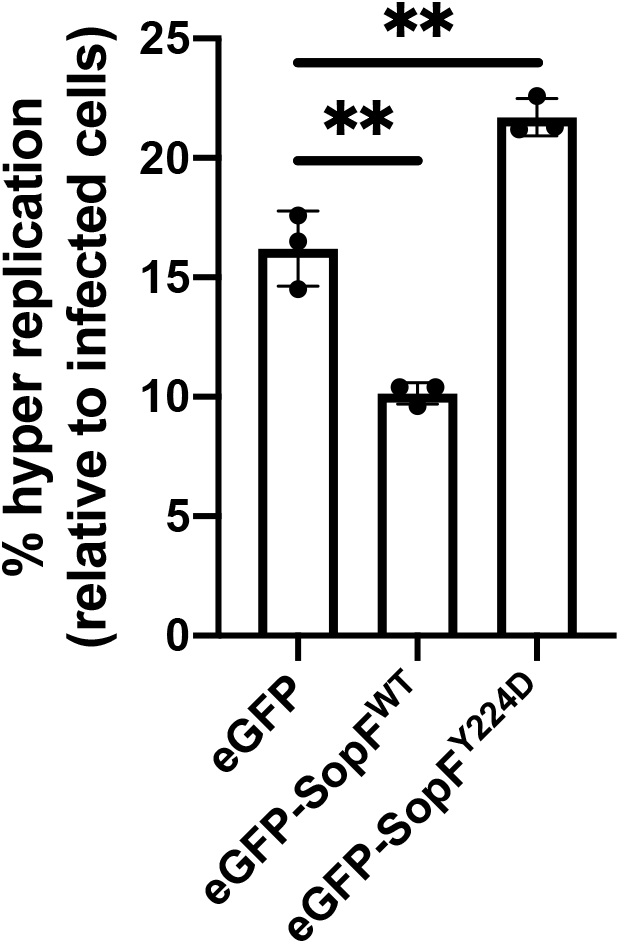
Ectopic SopF expression reduces cytosolic *S.* Typhimurium in HeLa cells. Quantification of the abundance of *S.* Typhimurium hyper replication in transgene-expressing infected HeLa cells at 6h pi using flow cytometry (n = 3). Significance cut-offs: ** = *P* < 0.01

**Figure S10.**
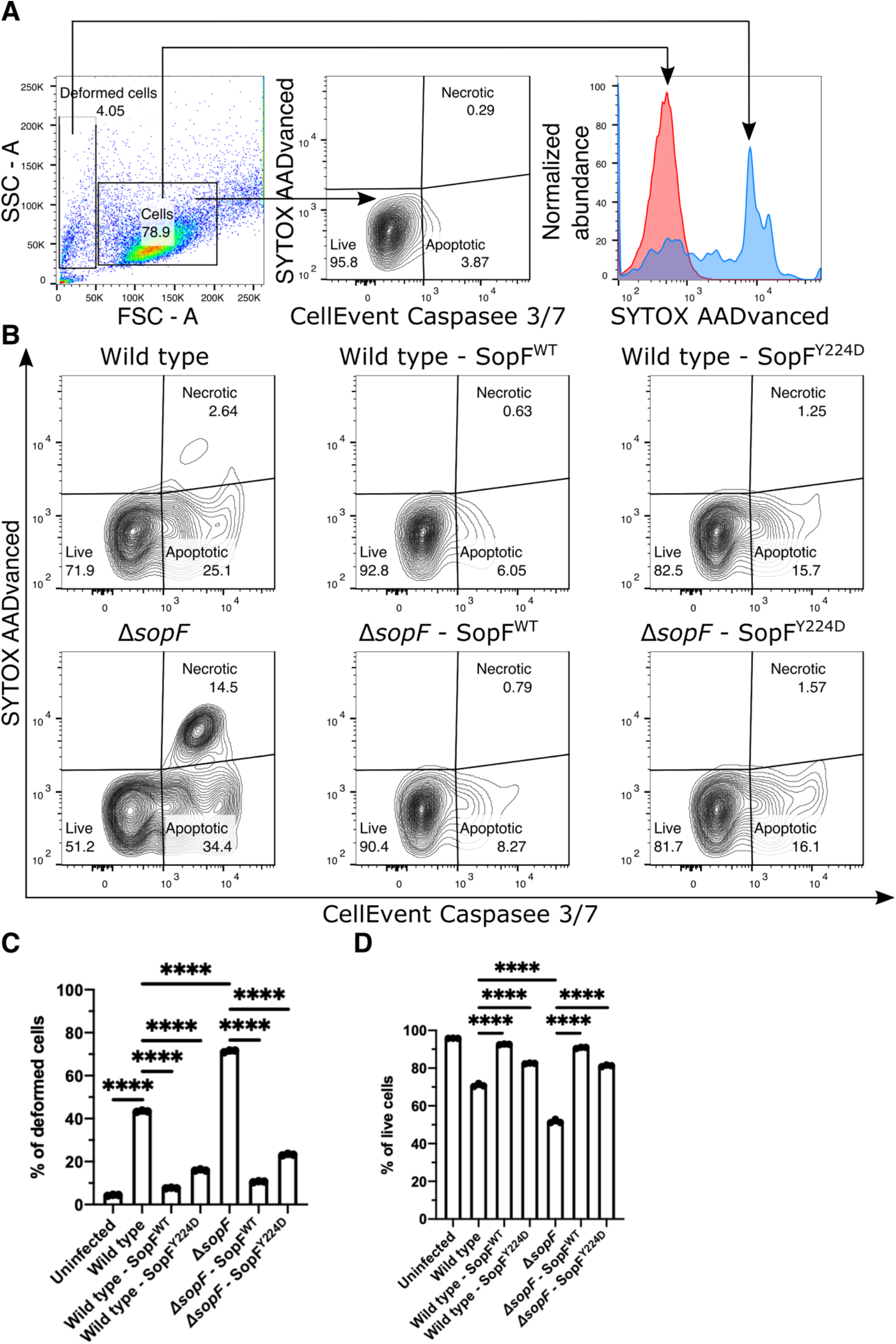
SopF suppresses host cell apoptotic and necrotic death. (A) Gating strategy for Caspase activity and cell death measurements. Uninfected or infected HeLa cells harvested at 6h pi were treated with CellEvent Caspase 3/7 and SYTOX AADvanced Dead Cell Stain and analyzed by flow cytometry. The “cells” and “deformed cells” were first gated from the total recorded events on the SSC-A vs FSC-A plot to remove cell debris. The “cells” were then visualized on the SYTOX AADvanced vs CellEvent Caspase-3/7 plot, where SYTOX^−^CellEvent^−^ were defined as live cells, SYTOX^−^CellEvent^+^ were defined as apoptotic cells and SYTOX^+^CellEvent^+^ were defined as necrotic cells. (B) Representative plots on the distribution of live, apoptotic and necrotic cells in various *S*. Typhimurium strains-infected HeLa cells at 6h pi (n = 3). (C) Quantification on the abundance of deformed cell population in HeLa cells at 6h pi (n = 3). (D) Quantification on the abundance of live cell in the gated “cells” at 6h pi (n = 3). Significance cut-offs: *** = *P* < 0.001 and **** = *P* < 0.0001.

**Figure S11.**
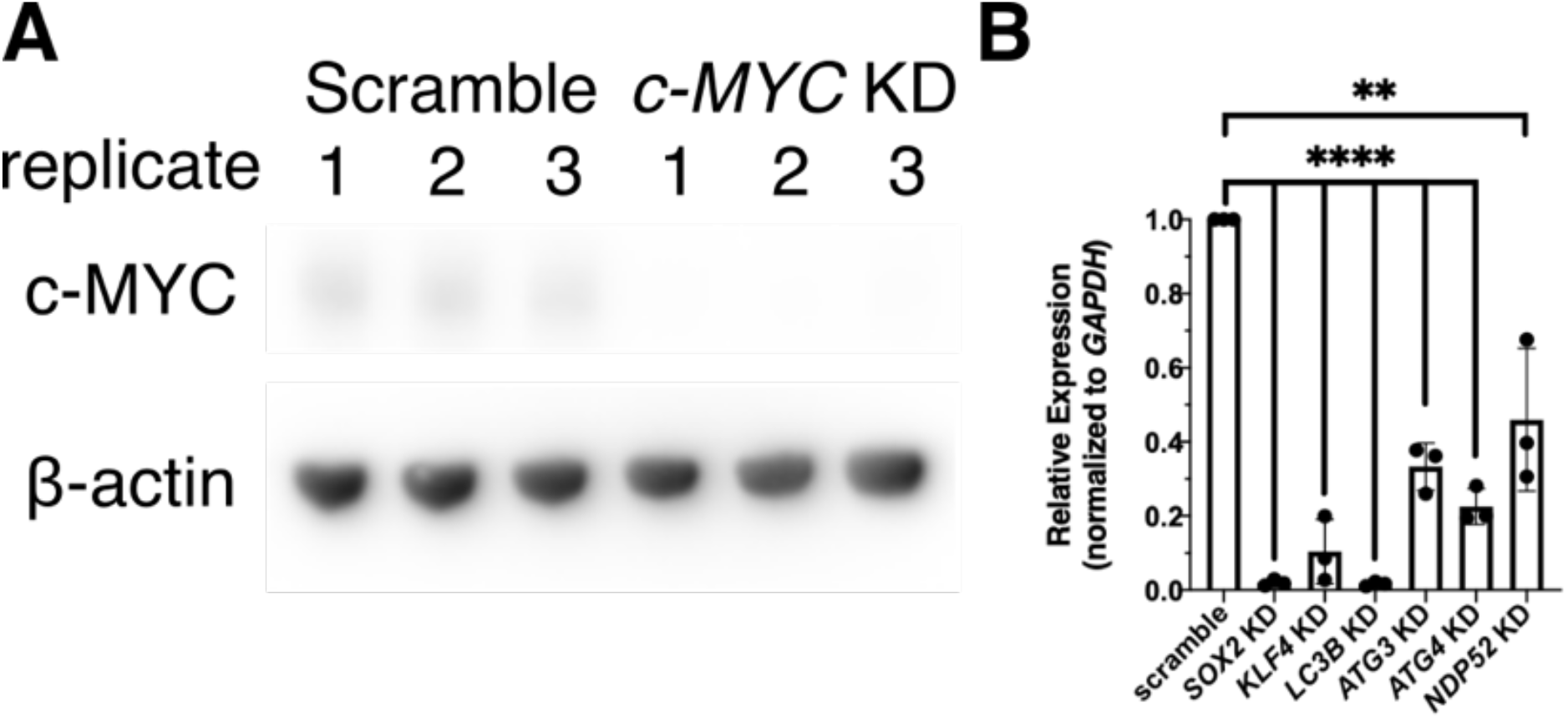
Evaluation of siRNA efficiency. (A) Evaluation on the siRNA knockdown efficiency of *c-MYC* using western blot (n = 3). (B) Quantification on the siRNA knockdown efficiency of *SOX2*, *KLF4*, *LC3B*, *ATG3*, *ATG4* and *NDP52* using qRT-PCR (n = 3). Significance cut-offs: ** = *P* < 0.01 and **** = *P* < 0.0001

**Table S1. Filtered candidates with >=2-folds enrichment**

In excel file

**Table S2. Filtered candidates with <2-folds enrichment**

In excel file

**Table S3.** List of bacteria strains.

| Name | Genotype | Plasmid | Reference |
| --- | --- | --- | --- |
| JL129 | wildtype | pSINA1.1 | Luk et. al. unpublished |
| JL177 | wildtype | pSINA1.7 | Luk et. al. unpublished |
| JL187 | $\Delta sopF$ | pSINA1.1 | This study |
| JL188 | $\Delta sopF$ | pSINA1.7 | This study |
| JL189 | $\Delta sopF$ | pSINA1.1 P<sub>BAD</sub> *sopF*<sup>WT</sup> | This study |
| JL190 | $\Delta sopF$ | pSINA1.1 P<sub>BAD</sub> *sopF*<sup>Y224D</sup> | This study |
| JL191 | $\Delta sopF$ | pBAD *sopF*<sup>WT</sup> | This study |
| JL192 | $\Delta sopF$ | pBAD *sopF*<sup>Y224D</sup> | This study |
| JL193 | wildtype | pSINA1.1 P<sub>BAD</sub> *sopF*<sup>WT</sup> | This study |
| JL194 | wildtype | pSINA1.1 P<sub>BAD</sub> *sopF*<sup>Y224D</sup> | This study |
| JL195 | wildtype | pBAD *sopF*<sup>WT</sup> | This study |
| JL196 | wildtype | pBAD *sopF*<sup>Y224D</sup> | This study |

**Table S4.** List of plasmids.

| Name | Description | Reference |
| --- | --- | --- |
| VSV-G | Encodes | Carette et al., 2009 |
| pAdvantage | Enhancement of retroviral production | Carette et al., 2009 |
| gag-pol | Encodes | Carette et al., 2009 |
| pGeneTrap | Encodes retroviral gene trap cassette | Carette et al., 2009 |
| pSINA1.1 | Detection of *S. Typhimurium* lifestyles | Luk et al. unpublished |
| pSINA1.7 | Detection of *S. Typhimurium* lifestyles | Luk et al. unpublished |
| pBAD *sopF*<sup>WT</sup> | Inducible expression of *sopF*<sup>WT</sup> | This study |
| pBAD *sopF*<sup>Y224D</sup> | Inducible expression of *sopF*<sup>Y224D</sup> | This study |
| pSINA1.1 P<sub>BAD</sub> *sopF*<sup>WT</sup> | Detection of *S. Typhimurium* lifestyles, complementation with *SopF*<sup>WT</sup> | This study |
| pSINA1.1 P<sub>BAD</sub> *sopF*<sup>Y224D</sup> | Detection of *S. Typhimurium* lifestyles, complementation with *SopF*<sup>Y224D</sup> | This study |
| pSINA1.7 P<sub>BAD</sub> *sopF*<sup>WT</sup> | Detection of *S. Typhimurium* lifestyles, complementation with *SopF*<sup>WT</sup> | This study |
| pSINA1.7 P<sub>BAD</sub> *sopF*<sup>Y224D</sup> | Detection of *S. Typhimurium* lifestyles, complementation with *SopF*<sup>Y224D</sup> | This study |
| peGFP-C3 | Negative control for ectopic expression in mammalian cells | Clontech (TaKaRa) |
| peGFP-*sopF*<sup>WT</sup> | Ectopic expression of *sopF*<sup>WT</sup> in mammalian cells | This study |
| peGFP-*sopF*<sup>Y224D</sup> | Ectopic expression of *sopF*<sup>Y224D</sup> in mammalian cells | This study |
| pBAD smURFP-HO-1 | Inducible expression of smURFP | Rodriguez et al., 2016 |

**Table S5.**
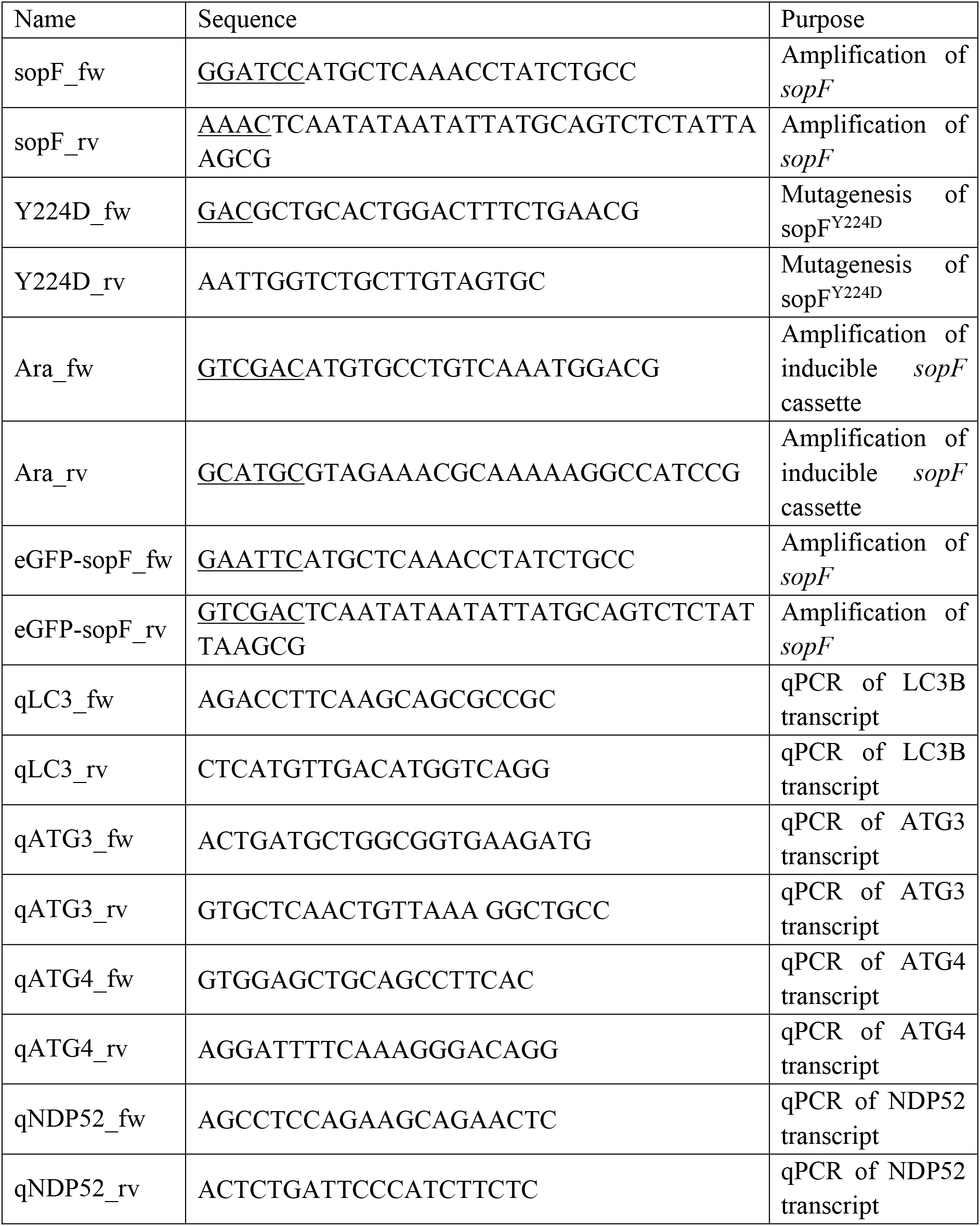
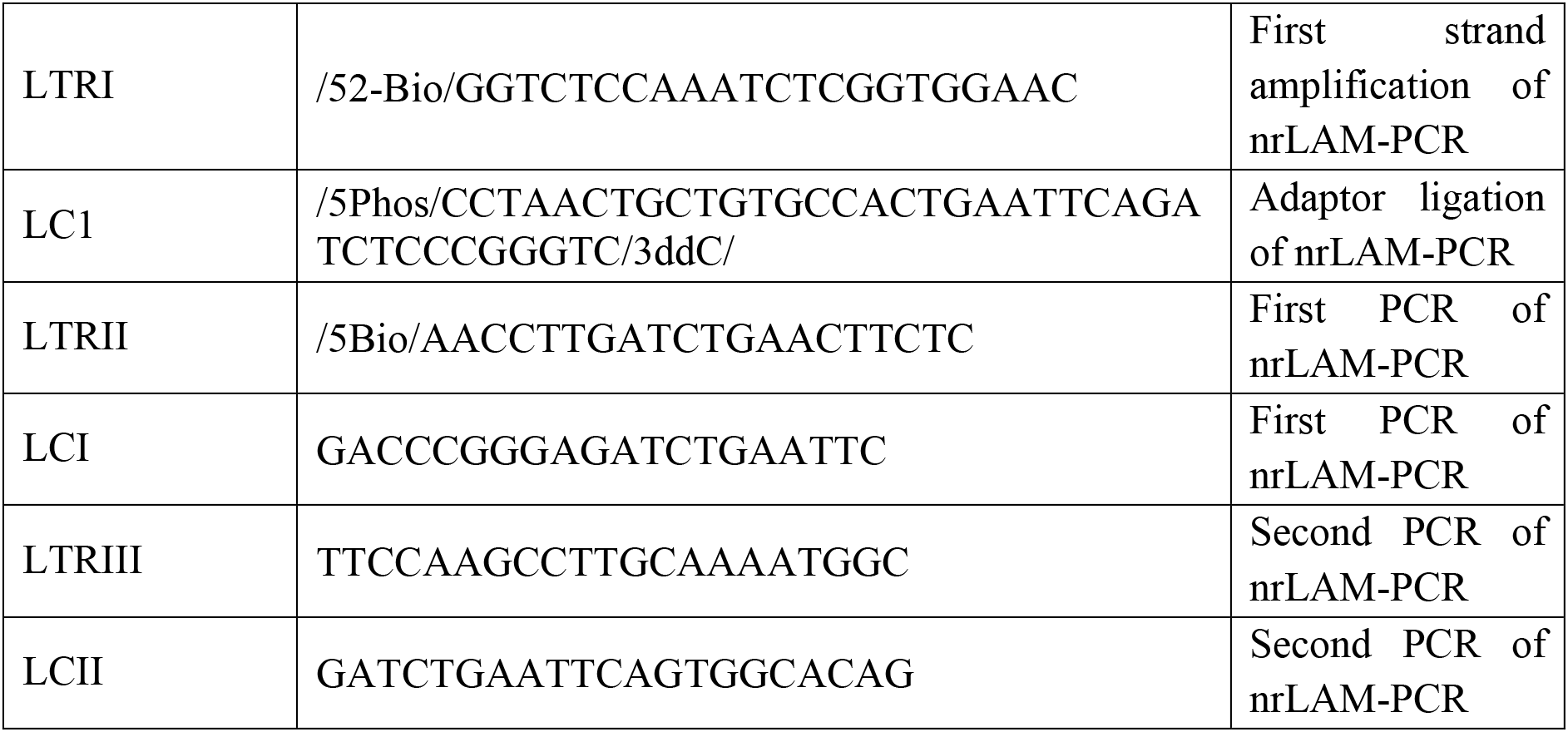
List of primers.

**Table S6.**
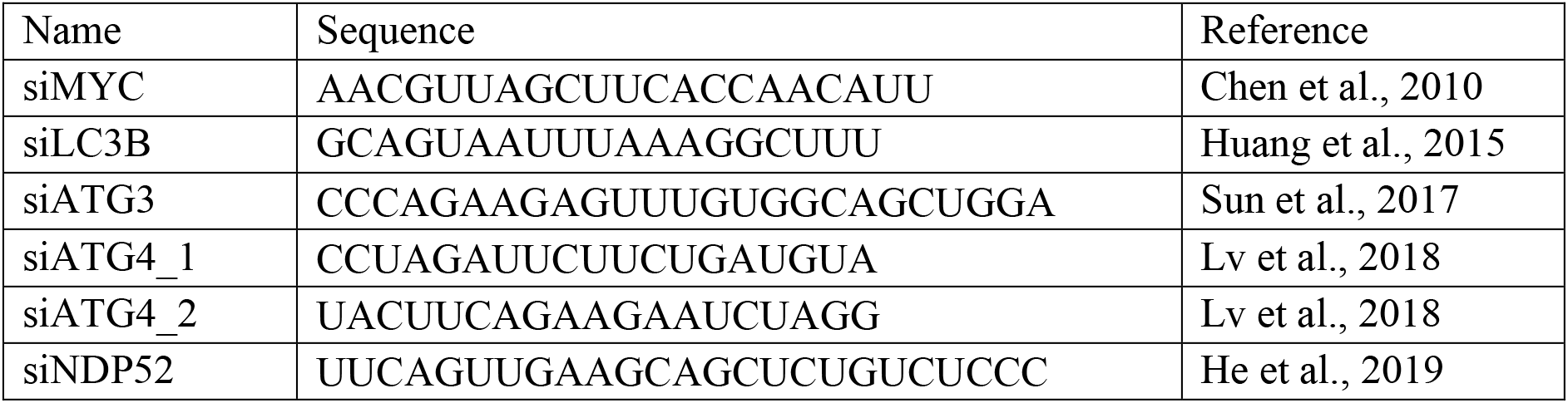
List of siRNA.

**Table S7.** List of reagents.

| Name of reagent | Concentration | Supplier | Reference |
| --- | --- | --- | --- |
| Anti-c-MYC antibody | 1:3000 | Abcam | Ab32072 |
| Anti-LC3 antibody | 1:200 | Proteintech | 14600-1-AP |
| Anti-Gelectin 3 antibody (M3/38) | 1:200 | SCBT | Sc-23938 |
| Monoclonal Anti-β-Actin antibody produced in mouse | 1:3000 | Sigma | A5316 |
| Goat anti-Rabbit IgG (H+L) Cross-Adsorbed Secondary Antibody, Alexa Fluor 405 | 1:200 | Invitrogen | A31556 |
| Goat anti-Rabbit IgG (H+L) Highly Cross-Adsorbed Secondary Antibody, Alexa Fluor 488 | 1:200 | Invitrogen | A11034 |
| Goat anti-Rat IgG (H+L) Cross-Adsorbed | 1:200 | Invitrogen | A10525 |
| Secondary Antibody, Cyanine5 |  |  |  |
| Goat Anti-Rabbit IgG (H + L)-HRP Conjugate | 1:3000 | Bio-Rad | 1706515 |
| Goat Anti-Mouse IgG (H + L)-HRP Conjugate | 1:3000 | Bio-Rad | 1706516 |

**Table S8.** Summary of NGS reads analysis.

|  |  | Control | Replicate 1 | Replicate 2 | Replicate 3 |
| --- | --- | --- | --- | --- | --- |
| Raw reads without adaptor |  | 3 982 389 | 780 852 | 722 999 | 2 908 085 |
| Merged reads |  | 3 680 138 | 686 722 | 642 104 | 2 680 051 |
| Mapped reads |  | 3 679 894 | 686 617 | 642 015 | 2 679 847 |
| Reads with $\geq 30$ nts<br>Genetrap sequence | | 1 137 870 | 52 945 | 266 877 | 641 677 |
| Annotated<br>reads | Sum | 1 137 870 | 52 945 | 266 877 | 641 677 |
|  | Exon | 76 019 | 5 828 | 7 919 | 21 047 |
|  | UTR | 125 | 275 | 3 439 | 6 531 |
|  | Intron | 542 125 | 28 635 | 109 509 | 331 029 |

